# Genomic variation and ancestry of *Helicobacter pylori* in the admixed population of Cabo Verde reveal host adaptation, limited host–pathogen ancestry concordance, and signatures of trans-Atlantic slave trade migrations

**DOI:** 10.64898/2026.06.05.730320

**Authors:** Timothy Archampong, Yi Ling Tam, Charlotte Davison, Isra Boughanmi, Sam Tallman, Milani Rodrigues, Neia Tavares, Kleyde Simone Santos, Francisco Barbosa Amado, Isabel Ramos, Antonieta Martins, Richard Haigh, Marco R. Oggioni, Rui Manuel Ferreira, Céu Figueiredo, Fabian Freund, Isabel Inês Araújo, Sandra Beleza

## Abstract

The genomic structure of the stomach-colonising, gastric cancer–associated bacterium *Helicobacter pylori* mirrors human evolutionary history, making it difficult to disentangle shared ancestry from true host–pathogen interactions driving disease risk between populations. Recently admixed populations offer opportunities for new coevolutionary trajectories between *H. pylori* and its host that may disrupt parallel evolution. Here, we analyse host *H. pylori* IgG, blood serum biomarkers, and paired host-bacterial genomic data from the general population (independent of dyspepsia symptoms) of Cabo Verde. Seropositivity is high (82.8%), as in neighbouring African populations. Serum biomarkers indicate that this unique cohort mainly comprises hosts with limited evidence of atrophic gastritis. Genomic analyses show that Cabo Verdean *H. pylori* derive European (hspSWeurope) and West African (hspAfrica1WAfrica) ancestry. The delineation of a reference dataset that includes new strains from Ghana, uncovered a distinct West– Central African cluster (hspAfrica1WCAfrica). This dataset allowed inference of within-Africa and within-Europe ancestries in Cabo Verdean and American strains that reflect historical migrations since European colonisation and the trans-Atlantic Slave Trade. European strains in Cabo Verde show additional population structure, including two low-diversity clusters. Approximate Bayesian Computation (ABC) inference indicates that the most common cluster represents a clonal expansion. Notably, these strains lack known virulence genes and show significant divergence from gastric cancer strains. The genomic signatures of this expanded group are compatible with founder effects and/or genetic adaptation, and new methods are needed to distinguish between these processes in highly recombining bacteria. Finally, host and bacterial ancestry are uncoupled, indicating disruption of ancestral parallel evolution. Cabo Verde, therefore, provides a unique natural model for studying host–pathogen interactions underlying susceptibility to *H. pylori*.

## Introduction

*Helicobacter pylori* is an obligate human-associated bacterium whose evolutionary history is intrinsically linked to humans’ history of migration since their origins in Africa 100,000 years ago (Falush et al., 2003; Linz et al., 2007; Moodley et al., 2012). Such codiversification has led to global geographic patterns of genetic diversity in *H. pylori* that are congruent with those of their host (Suerbaum and Josenhans 2007). As a result, population structure in *H. pylori* can be partitioned into major populations (“hp”) and subpopulations (“hsp”) with distinct geographic distributions, and patterns of genetic variation within these groups have been used to infer the demographic history of their host (Wirth et al., 2004; Moodley et al., 2021; Suzuki et al., 2022). Recently, the *Helicobacter pylori* Genome Project (HpGP) has generated over 1000 *H. pylori* high-quality genomes from 50 countries to refine the bacterium’s worldwide population structure and evolution, and which comprises a valuable reference dataset to evaluate population history and ancestry of bacterial populations across different geographical regions (Thorell et al., 2023).

Despite being a frequent commensal coloniser of the gastric mucosa, *H. pylori* also exhibit great pathogenic potential and are a major risk factor for gastric cancer (de Martel et al., 2020). Several genes in *H. pylori* have been associated with increased risk of gastric cancer such as the *cagA* (cytotoxin-associated gene) pathogenicity island (cagPAI), *vacA* (vacuolating cytotoxin), and *babA* (Lewis blood group antigen-binding adhesin)(Yamaoka 2010; Linz et al., 2024). The presence of one or more of these genes does not however necessarily lead to disease (Linz et al., 2024). In fact, despite commonly possessing virulence genes, the majority of *H. pylori* colonisations result in lifelong asymptomatic carriage; only 1-5% of the *H. pylori*-positive individuals develop gastric cancer, depending on ethnicity and environmental factors (Malfertheiner et al., 2023). This indicates that disease risk in *H. pylori* infection is driven by complex combinations of bacterial virulence, host genetic susceptibility, and environmental diversity.

Parallel evolution and correlated population structure between *H. pylori* and humans makes it difficult to distinguish which cooccurrence of human-bacterium variants are due to parallel ancestral lineages and which are responsible for real biological interactions underlying differences in susceptibility across host populations. One way to overcome this problem is to analyse human-*H. pylori* interactions in recently admixed populations. Admixture brings together diverse genetic backgrounds from both the host and the microbe, thereby creating novel host-bacterium combinations that are absent in non-admixed populations and disrupting the ancestral genetic correlation between *H. pylori* and its host. For instance, although the genetic ancestry of *H. pylori* was associated with the genetic ancestry of their host in patients from two Colombian communities, African strains were more virulent in hosts with high Native American ancestry, explaining the difference in gastric cancer incidence between those communities (Kodaman et al., 2014).

Recently admixed populations with different admixture histories comprise novel opportunities for new evolutionary and coevolutionary trajectories between *H. pylori* and its host (Thorell et al., 2017; Muñoz-Ramirez et al., 2021). For instance, “New World” strains from North, Central and South America showed highest divergence from “Old World” strains at virulence genes; this variation might contribute to higher virulence potential of the American strains. The disruption of previous parallel ancestry in these populations seems to lead to more pathogenic outcomes, akin to that seen in Kodaman et al. (2014).

Here, we evaluate *H. pylori* genomic variation, population structure, and host-*H. pylori* codiversification trajectories that have arisen in Cabo Verde, an Altantic archipelago off the coast of West Africa. Colonized by the Portuguese in the mid-15th century, the Cabo Verdean population derive mixed ancestry from Europe and West Africa (Beleza et al., 2012; Beleza et al., 2013; Korunes et al., 2022; Laurent et al., 2023). We hypothesized that the unique population history and patterns of admixture will have broken down the correlation between host and *H. pylori* ancestry and genetic variation, and that Cabo Verde can, therefore, be considered as a model to evaluate host-*H. pylori* interactions underlying disease risk.

At the outset of this study, no epidemiological surveillance of *H. pylori* infection had been conducted in the islands. Therefore, we recruited volunteers from the general population, independently of the presence of dyspeptic symptoms, and collected paired host and *H. pylori* genomic data. This integrated dataset enabled us to: (i) conduct an unbiased epidemiological assessment of *H. pylori* colonisation in Cabo Verde; (ii) assess *H. pylori* genetic variation associated with human migrations during the Trans-Atlantic Slave trade; (iii) characterise the between-host and population-level *H. pylori* genetic variation, ancestry and evolution in Cabo Verde; and iv) evaluate the relationship between host and *H. pylori* ancestries across the islands and assess the extent to which Cabo Verde constitutes a natural model for disentangling host–microbe coevolution and disease-related interactions.

## Results

### *H. pylori* epidemiology and association with gastric inflammation indicators in Cabo Verde

In total, 876 Cabo Verdeans of the general population (regardless of dyspeptic symptoms) were evaluated for of *H. pylori* IgG in blood serum (Table 1). There is an over-representation of women in the cohort (73% of the sample); the age ranged from 23 to 93 years old (mean ± SD= 43 ± 13.2). While most individuals (65%) were classified as being from Santiago, our two-generation island descent criterion provided a comparable dataset for the remaining islands (Table 1). The cohort contained related individuals: 51 pairs/trios of first-degree relatives (either parents/offspring or sibling pairs/trios), three pairs/trios of second-degree relationship (one grandparent/grandchildren trio, one half-siblings trio, and one aunt/niece pair), and 24 couples. There was no difference in seropositivity between related and unrelated groups. Nonetheless, statistical analyses excluded related individuals.

**Table 1.**
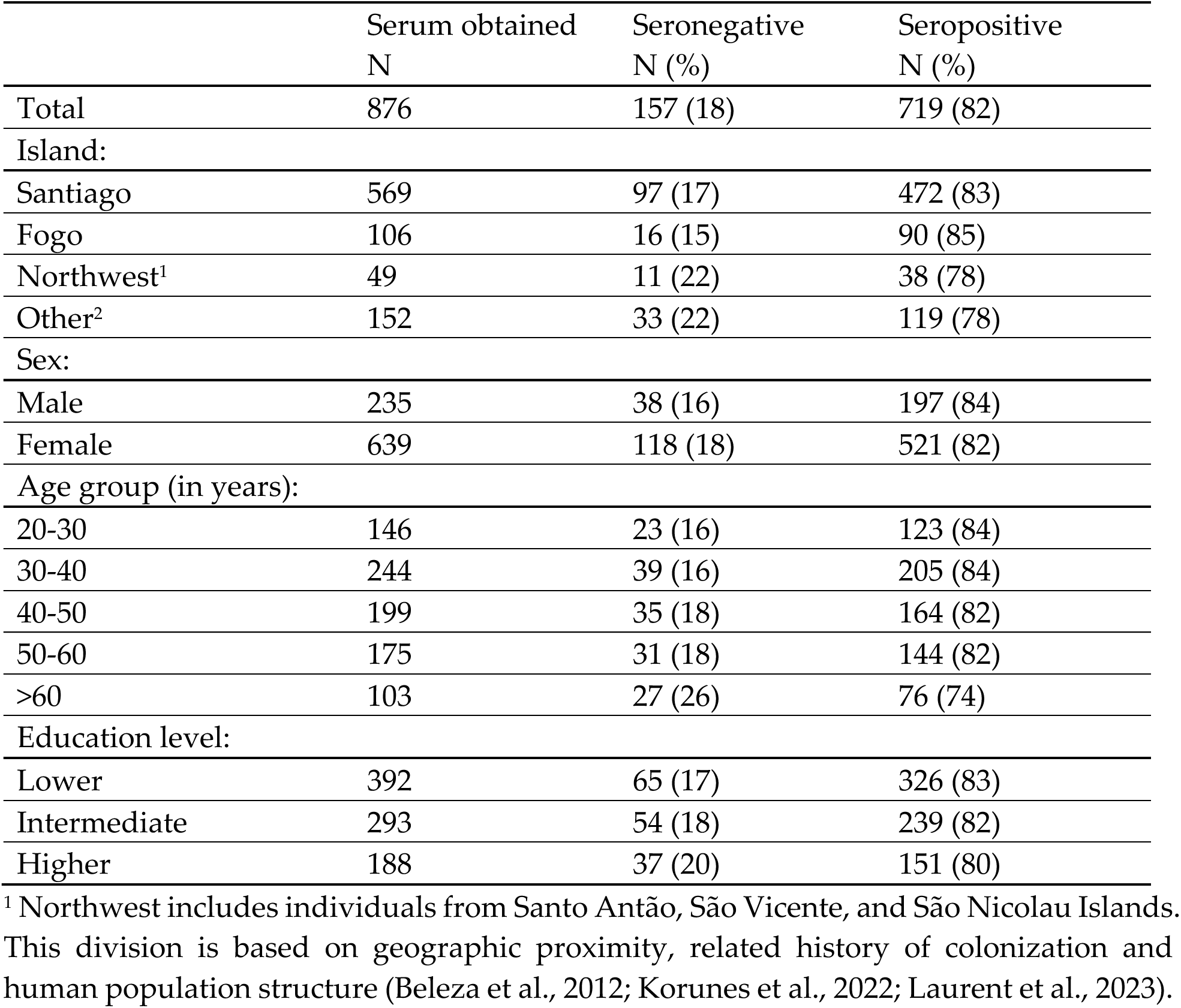
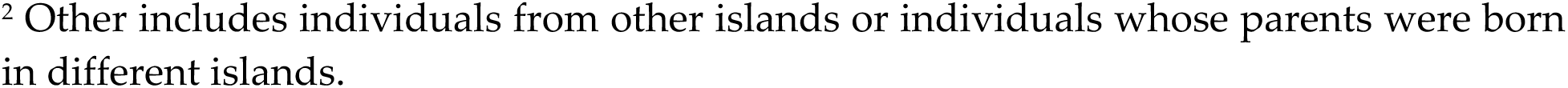
with 1 supplement Socio-demographic characteristics of participants in the epidemiological survey, in total sample and stratified by *H. pylori* serostatus.

|  | Serum obtained<br>N | Seronegative<br>N (%) | Seropositive<br>N (%) |
| --- | --- | --- | --- |
| Total | 876 | 157 (18) | 719 (82) |
| Island: |  |  |  |
| Santiago | 569 | 97 (17) | 472 (83) |
| Fogo | 106 | 16 (15) | 90 (85) |
| Northwest<sup>1</sup> | 49 | 11 (22) | 38 (78) |
| Other<sup>2</sup> | 152 | 33 (22) | 119 (78) |
| Sex: |  |  |  |
| Male | 235 | 38 (16) | 197 (84) |
| Female | 639 | 118 (18) | 521 (82) |
| Age group (in years): |  |  |  |
| 20-30 | 146 | 23 (16) | 123 (84) |
| 30-40 | 244 | 39 (16) | 205 (84) |
| 40-50 | 199 | 35 (18) | 164 (82) |
| 50-60 | 175 | 31 (18) | 144 (82) |
| >60 | 103 | 27 (26) | 76 (74) |
| Education level: |  |  |  |
| Lower | 392 | 65 (17) | 326 (83) |
| Intermediate | 293 | 54 (18) | 239 (82) |
| Higher | 188 | 37 (20) | 151 (80) |

<sup>1</sup> Northwest includes individuals from Santo Antão, São Vicente, and São Nicolau Islands. This division is based on geographic proximity, related history of colonization and human population structure (Beleza et al., 2012; Korunes et al., 2022; Laurent et al., 2023).
<sup>2</sup> Other includes individuals from other islands or individuals whose parents were born in different islands.

There is no association between *H. pylori* serostatus and island group (or by island), sex or education level. Individuals with ages >60 years were significantly less seropositive than the other age groups (Odds Ratio = 0.5; 95% CI: 0.2-0.9; p-value= 0.035). This is associated with lower IgG concentration specially for individuals >70 years old (>70 years, n=29, regression coefficient, coeff (S.E.) = -16.6 (8.1) µg/L, p-value= 0.04; Table 1-figure supplement 1A). This suggests that age-related decline in antibody levels or sero-reversion contributes to a lower seropositivity in older individuals. The overall seropositivity in Cabo Verde (calculated for individuals <70 years old) is 82.8% (95%CI: 80.0-85.6%). It should be noted that seropositivity reflects both active and past infection. Therefore, the prevalence of active infection may be lower than the observed seropositivity rate (see Discussion).

Evaluation of the serum levels of the pepsinogen I (PgI), pepsinogen II (PgII) and the PgI/PgII ratio allowed us to indirectly assess gastric inflammation (individuals with active gastritis have elevated PgII levels) and gastric mucosal atrophy (PgI < 30 µg/L and/or PgI/PgII ratio < 3 indicate increased risk of atrophic gastritis) of the individuals (Table 1 – figure supplement 1) (Di Mario et al., 2006; Telaranta-Keerie et al., 2010; Kitamura et al., 2015). Although PGI and PGII levels are significantly elevated in seropositive individuals, 90% and 80% of the seropositive individuals have PgI and PgII levels in the same range as seronegative individuals (Table 1 – figure supplement 1B and 1C). PgI/PgII < 3 were found in 19 individuals (2.2% of the total cohort, n=876) and were associated both with lower levels of PgI (Kruskal-Wallis test, p-value= 9.2×10^-9^; all except two individuals had PgI < 30 µg/L) and higher levels of PgII (Kruskal-Wallis test, p-value= 0.001). Overall, PgI/PgII ratio < 3 was associated with *H. pylori* seropositivity (1.7% (n=12) of seropositive individuals; chi-square = 4.8, 1 degree of freedom, p-value= 0.03; Table 1 – figure supplement 1D).

Taken together, the serological data provide limited evidence of gastric mucosal atrophy within the cohort, despite the high prevalence of seropositivity.

### Subcontinental ancestry in Cabo Verdean and American *H. pylori* reflect historical host migrations since the Trans-Atlantic slave trade

We sequenced 550 *H. pylori* colonies isolated from 178 hosts (range= [1-33]; mean = 3 per host) and calculated the number of pairwise Single Nucleotide Variants (SNVs) across all core genomes. Relatedness Estimation in Admixed Populations (REAP) analysis (Thornton et al., 2012) identified seven second-degree and seven first-degree relative pairs among the hosts, corresponding to a total of 28 related individuals. For all related pairs except one, the number of pairwise SNVs (range = [2,400-35,938]) overlapped with those observed for unrelated hosts (range = [983-40,392]; except for one mother/daughter pair whose isolates differed by 240 SNVs). In 35 hosts with more than one isolate sequenced, within-host pairwise SNV differences also overlapped with the range of observed between unrelated hosts, indicating multiple colonization in ∼20% of the individuals. Given this, we assembled a dataset of 220 unrelated *H. pylori* isolates (with >1000 pairwise SNV differences) from 177 individuals from Cabo Verde, along with one *H. pylori* isolate from Senegal (full participant flow is in Figure 1 – figure supplement 1). We then ran haplotype-based ChromoPainter/fineSTRUCTURE (Lawson et al., 2012) and genotype-based ADMIXTURE (Alexander et al., 2009) analyses with a comprehensive worldwide isolate dataset (including those from but not restricted to the HpGP dataset; Figure 1 – source data 1), and the Cabo Verdean isolates to further investigate population structure and ancestry in the *H. pylori* populations resulting from historical migrations from European and African continents (Figure 1; Figure 1 – figure supplement 2 – 9).

**Figure 1.**
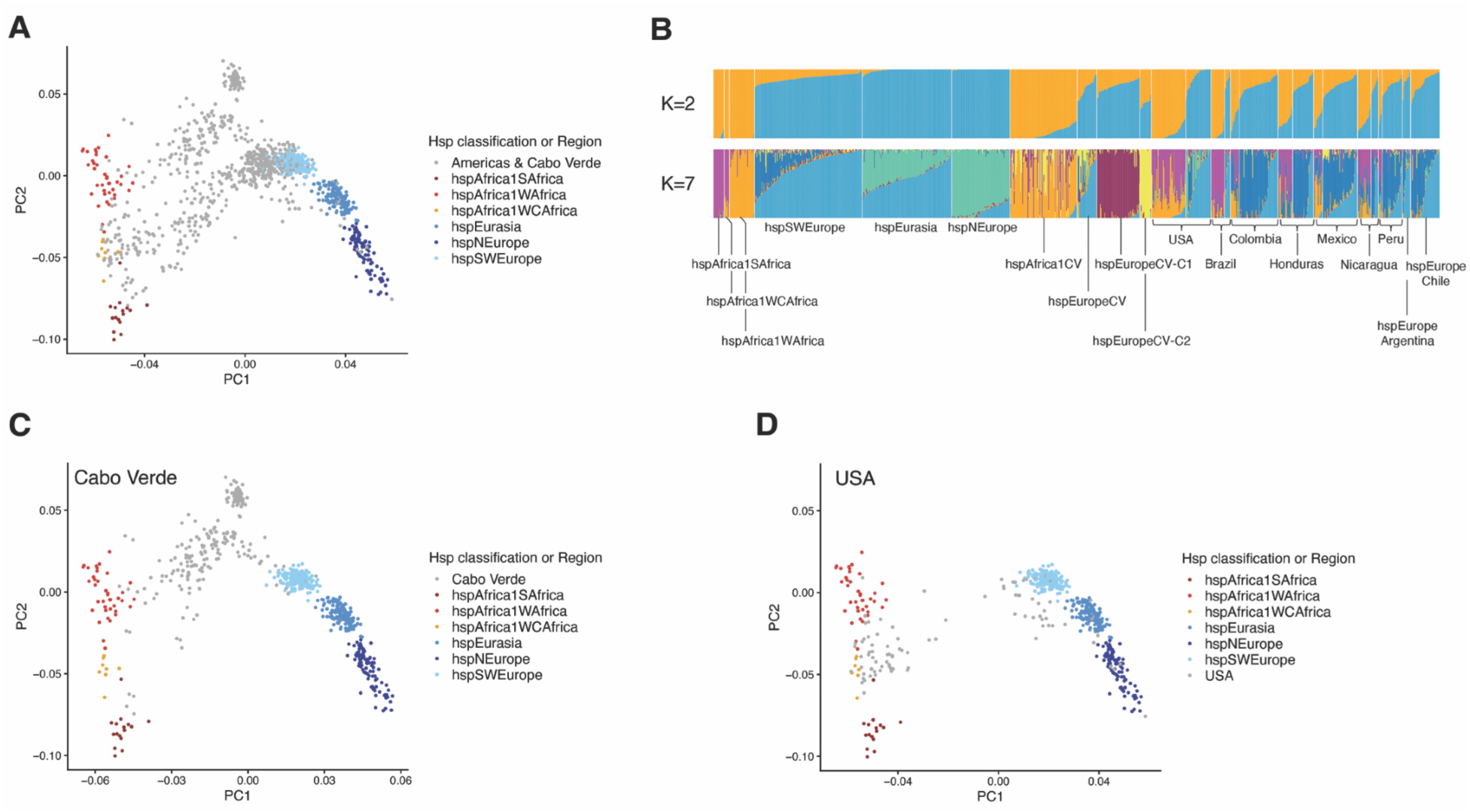
with 10 supplements **Population structure and ancestry of *H. pylori* strains from Europe, African, the Americas and Cabo Verde.** (A) Coancestry PCA analysis of 1165 *H. pylor*i strains from Europe, Africa, multiple American populations and Cabo Verde. (B) Unsupervised ADMIXTURE Clustering at K=2 and K=7. Only European and African population groups with *n* > 10 are included. Cabo Verdean samples are subdivided according to within-population structure (see Figure 2; hspEuropeCV-C1 - hspEuropeCV-Clonal1; hspEuropeCV-C2 - hspEuropeCV-Clonal2). American populations are assigned to either hspEurope or hspAfrica1 based on their proportion of European and African ancestry. Accordingly, strains from the USA, Brazil, Colombia, Honduras, Mexico, Nicaragua, and Peru are divided in, from left to right, hspAfrica1USA, hspEuropeUSA, hspAfrica1Brazil, hspEuropeBrazil, hspAfrica1Colombia, hspEuropeColombia, hspAfrica1Honduras, hspEuropeHonduras, hspAfrica1Mexico, hspEuropeMexico, hspAfrica1Nicaragua, hspEuropeNicaragua, hspAfrica1Peru, hspEuropePeru. (C) Same PCA as in (A), but showing only the Cabo Verdean sample. (D) Same PCA as in (A), but showing only the USA population. Sample sizes are in Figure 1-source data 1.

We first inferred the continental ancestry of Cabo Verdean *H. pylori* using a subset of worldwide strains (Figure 1-source data 1; Figure 1-supplement figure 2). Cabo Verdean strains were found to spread between African and European populations in principal component analysis (PCA) of the coancestry matrix obtained with fineSTRUCTURE. There was no detectable genetic contribution from hpAmerind, hspEAsia, hpAsia2 and hpAfrica2 to the Cabo Verdean bacterial population and, therefore, these reference strains were, excluded from further analyses.

Next, to define geographic-localised reference groups, we considered as European and African references only those isolates from Europe or Africa whose ancestry was determined by PCA, k-means clustering and fineSTRUCTURE to be from their respective continents (Figure 1-supplement figure 3). In a preliminary fineSTRUCTURE analysis with Cabo Verdean and American isolates, African isolates from Sudan, Cameroon and Nigeria were found to be outlying in PC2 of the coancestry PCA and were, therefore, removed from subsequent analyses (Figure 1-supplement figure 4). The final analyses were performed with a total of 1165 isolates and 37,480 core SNVs with complete genotype data (Figure 1; Figure 1 - figure supplement 5-9 and Figure 1 – source data 1).

In the coancestry PCA resulting from this fineSTRUCTURE analysis, the European reference samples formed a cline of three clusters on PC2 (Figure 1A), consistent with previous findings from the HpGP (Thorell et al., 2023). Accordingly, they were classified into hspNEurope, hspEurasia and hspSWEurope, following the HpGP classification scheme. This dataset includes 47 new genomes from Portugal from our own collection, all of which were classified as hspSWEurope.

In contrast, the addition of four new genomes from Ghana into the African reference dataset led to a new visible cline in PC2 (Figure 1A). These new Ghanaian isolates formed a new cluster together with three isolates from Ghana and Nigeria which have been previously classified as hspAfrica1NAmerica by the HpGP (Thorell et al., 2023). fineSTRUCTURE MCMC tree assignments inferred from the full coancestry matrix further support the existence of a distinct clade separate from hspAfrica1WAfrica and hspAfrica1SAfrican (Figure 1-figure supplement 5). This clustering pattern persisted in ChromoPainter/fineSTRUCTURE analyses without the four new Ghanaian isolates and when we downsampled the other hspAfrica1 clusters (Figure1-figure supplements 6-7). This indicates that the detection of the new cluster was not driven by uneven sample sizes among the African clusters. As this cluster is positioned between the previously described hspAfrica1WAfrica and hspAfrica1SAfrica and, in accordance with its geographic location in Africa, we refer to it here as hspAfrica1WestCentralAfrica (hspAfrica1WCAfrica).

These fineSTRUCTURE and ADMIXTURE (K=2-7) analyses allowed us to detect, for the first time, differences in subcontinental European and African ancestry among *H. pylori* genomes from Cabo Verde and 14 American countries (Figure 1; Figure 1-figure supplement Supplementary 8-9). On the African side, Cabo Verdean isolates traced their ancestry predominantly to hspAfrica1WAfrica, whereas most of the American isolates traced their ancestry to hspAfrica1WCAfrica (e.g. USA, Colombia, Dominican Republic, Nicaragua, Peru; with most of these having previously been classified as hspAfrica1NAmerica, Figure 1 – source data 1), or to hspAfrica1WCAfrica and hspAfrica1SAfrica (e.g. Brazil, Honduras). On the European side, most of the American and Cabo Verdean isolates appear to trace their ancestry to hspSWEurope, although some USA, Argentinian, and Brazilians isolates are also found to have ancestry to hspEurasia and hspNEurope.

Due to sample size for most American *H. pylori* populations being limited, it is not possible to draw definitive conclusions about their overall ancestral origins; however, it is noteworthy that the observed patterns align with the subcontinental African and European ancestry observed for their human hosts (Ongaro et al., 2019; Micheletti et al., 2020; Laurent et al., 2023).

### There is additional population structure in Cabo Verdean *H. pylori*

The FineSTRUCTURE and ADMIXTURE K=7 analyses reveal additional population structure in Cabo Verdean *H. pylori* that is not solely explained by ancestry (Figure 1A-C; Figure 1-figure supplement 5). PCA and phylogenetic analyses of Cabo Verdean *H. pylori* core genomes disclose four main clusters or clades (Figure 2A-B): two clusters closely related with hspAfrica1 and hspSWEurope (Figure 2 - figure supplement 1), and two additional clusters of isolates that form short-branched, star-like phylogenetic clades. Individual ancestries estimated using SOURCEFIND (Chacon-Duque et al., 2018) with hspSWEurope and hspAfrica1WAfrica as surrogates and each isolate as a target were congruent with ADMIXTURE K=2 (Spearman correlation = 0.88; Figure 1b and Figure 2 - figure supplement 2). This analysis confirmed a major European ancestry component in these two unique Cabo Verdean groups. Estimation of population level ancestries allowing for self-copy with SOURCEFIND show that, unlike the references and other coexisting Cabo Verdean population groups, each of the two unique European-derived groups copy more haplotypes with each other that with any other population group (Figure 2C; Figure 2 – source data 1). In addition, the folded synonymous (neutral) site-frequency spectrum (_f_SFS, distribution minor-allele absolute frequency at synonymous variant sites) of these unique groups are skewed towards rare variation in comparison to the other coexisting Cabo Verdean population groups, and their genetic diversity is considerably lower (Figure 2D; Figure 2 - figure supplement 3). Collectively and in accordance with the definition of *H. pylori* clonal groups in previous studies (Thorell et al., 2023), these findings support that these two groups are comprised of closely related isolates resulting from recent clonal expansions. Following the nomenclature rules established by the HpGP but considering only continental-level ancestry of the isolates, the four Cabo Verdean *H. pylori* populations were termed hspAfrica1CaboVerde (n=103), hspEuropeCaboVerde (n=31), hspEuropeCaboVerde-Clonal1 (n=68) and hspEuropeCaboVerde-Clonal2 (n=19) (Figure 2).

**Figure 2.**
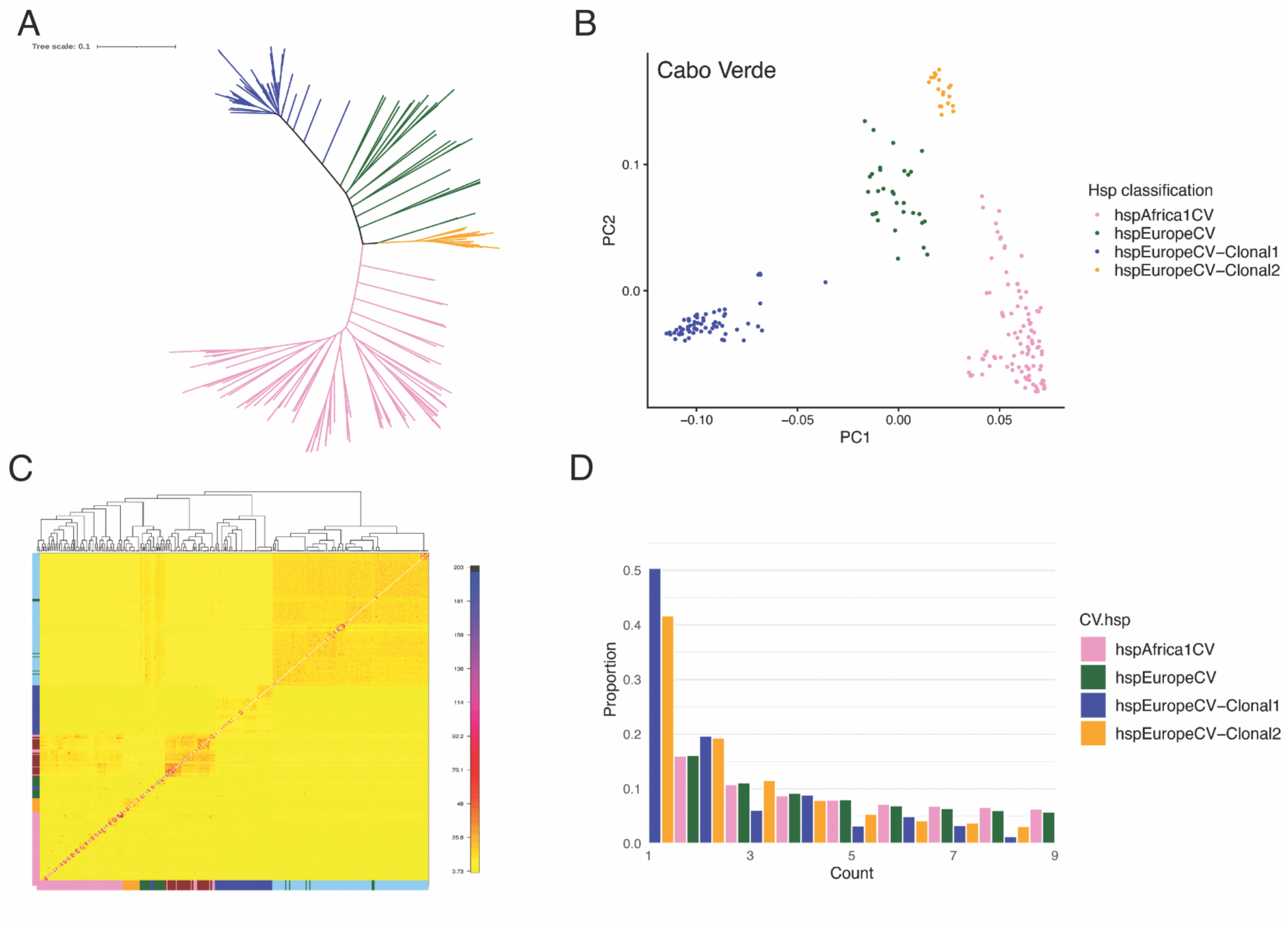
with 6 supplements **Population Structure of *H. pylori* strain in Cabo Verde.** (A) Maximum likelihood unrooted phylogenetic tree of Cabo Verdean strains. Colour code is in (B) and (D). (B) PCA of Cabo Verdean strains. (C) ChromoPainter co-ancestry matrix with each cell indicating the proportion of DNA chunks that each genome copies from all other genomes (see Figure 2 – data source 1 for SOURCEFIND self- and between group copy fraction estimates). The colour bar on the left is repeated in the bottom of the matrix and denotes the Cabo Verdean population groups (with colour code in B and (D)), hspSWEurope (light blue) and hspAfrica1WAfrica (red). (D) Folded site frequency spectrum (_f_SFS) of synonymous variants in Cabo Verdean population groups. To get a comparable result, the _f_SFS was calculated based on the same sample size for all population groups (n=19, as in hspEuropeCV-Clonal2). Sample sizes: hpsAfrica1CV, n=103; hspEuropeCV, n=31; hspEuropeCV-Clonal1, n=68; hspEuropeCV-Clonal2, n=19.

### Demographic inference supports increased population growth of hspEuropeCV*-*Clonal1 in Cabo Verde

We further investigated the demographic history of Cabo Verdean populations using the approximated Bayesian computation (ABC) approach based on the coalescent model from Davison et al. ((Davison et al., 2024); see methods), to infer a population growth model (exponential vs. no growth) and growth rate that best explain observed genetic variation in each Cabo Verdean population. Posterior predictive checks show that sample sizes do not affect model selection and only affect the growth rate estimation slightly (Figure 2 - figure supplement 4). Although growth rates have wide confidence estimates, our approach shows support for meaningful exponential population growth in hspEuropeCV-Clonal1 (growth rate estimate=13.7; 95% CI=[6.2,20.3]), followed by weaker growth in hspEuropeCV (growth rate estimate=1.1; 95% CI=[0.5,2]), and minimal growth in hspAfrica1CV (growth rate estimate =0.2; 95% CI=[0.1,0.5]). Due to limited sample size, we did not attempt to model the genetic diversity of hspEuropeCV-Clonal2 or infer its population growth rate.

Population expansion signatures due to an excess of rare variants is common in bacteria and other microorganisms (Lapierre et al., 2016; Davison et al., 2024; Goldberg 2026). Consistent with this, posterior predictive checks for Tajima’s D support a Kingman coalescence with exponential growth model for hspEuropeCV-Clonal1 (Figure 2 - figure supplement 4). However, alternative explanatory explanations cannot be excluded. In particular, the observed genealogical patterns and site-frequency spectrum may be also consistent with multiple-merger genealogies, which arise when reproductive success is highly skewed and a small number of lineages contribute disproportionately to the next generation (Menardo et al., 2021; Goldberg 2026). In bacterial populations, this type of genealogy can emerge through strong founder effects, transmission bottlenecks and other forms of genetic drift (Thorell et al., 2017; Menardo et al., 2021), as well as positive selection (Lapierre et al., 2016). Resolving the relative contributions of drift and selection to observed genetic variation will require a statistical framework that jointly evaluates the goodness-of-fit of inferred genealogies and performs model selection between competing evolutionary models. These approaches will need to account for high recombination to be applicable to hspEuropeCV-Clonal 1 and other *H. pylori* populations.

### Clonal expansion hspEuropeCV-Clonal1 is genetically differentiated from gastric cancer strains

To identify and characterize local adaptations within the hspEuropeCV-Clonal1 and hspEuropeCV-Clonal2 lineages, we calculated the Population Branch Statistic (PBS) for each core-genome SNV and accessory gene (Yi et al., 2010). PBS measures hspEuropeCV-clonal lineage-specific allele-frequency changes at a given locus relative to the divergence from hspEuropeCV and hspAfrica1CV, effectively identifying loci that have experienced rapid evolution since the divergence of these Cabo Verdean clonal groups.

Fold enrichment analysis of functional annotations for the most highly differentiated genes assigned based on eggNOG ortholog annotations (Huerta-Cepas et al., 2019) revealed a significant enrichment of core and accessory genes associated with cell wall and cell membrane biogenesis (COG category M) within hspEuropeCV-Clonal1(Figure 3A; Figure 3 – source data 1). Due to limited sample size, hspEuropeCV-Clonal2 did not show significant enrichment of any core or accessory COG categories (Figure 3 – source data 2).

**Figure 3.**
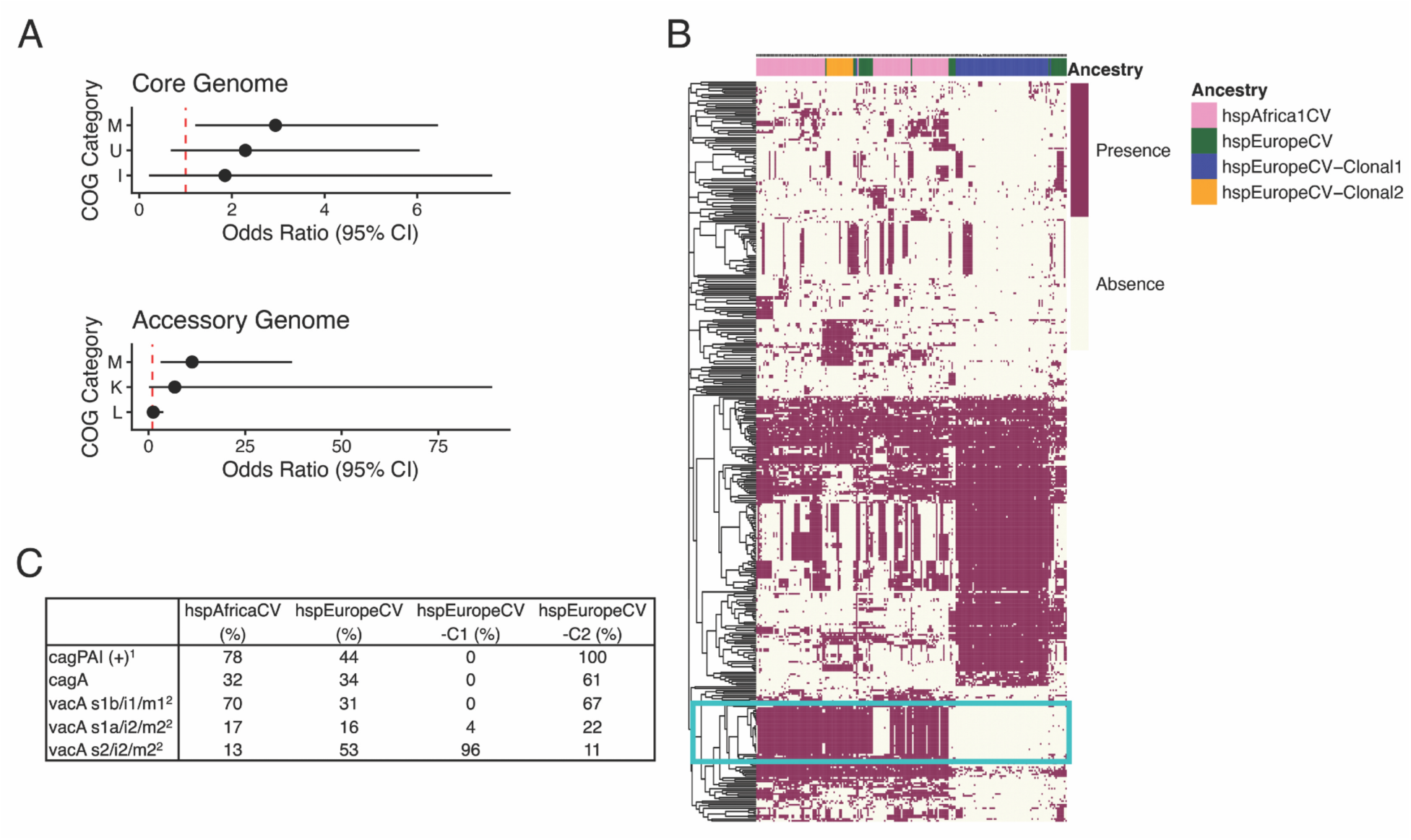
with 5 supplements **Genomic composition of Cabo Verdean *H. pylori* population groups**. (**A)** Forest plots showing odds ratios and 95% confidence intervals for the top three genomic annotations in the core and accessory genomes, derived from a fold enrichment analysis of hspEuropeCV–Clonal1. COG categories: M - Cell wall/membrane/envelope biogenesis; U - Intracellular trafficking, secretion, and vesicular transport; I - Lipid transport and metabolism; K-Transcription; L-Replication, Recombination and Repair. Core genetic variants were called via mapping to the reference sequence 26695 and correspond to those present in ≥95% of the total sample. **(B)** Hierarchical clustering based on the accessory genomes’ presence (burgundy)/absence (white) for the whole Cabo Verdean sample. Strains are coloured based on their population group on the top of the chart. Boxed region corresponds to *cagPAI.* Accessory genes were obtained through pangenome alignment and correspond to genes present in 5-95% of the total sample. **(C)** Frequency of active *cagPAI* (*cagPAI* (+)), *cagA*, and *vacA* genotypes in Cabo Verdean *H. pylori* population groups. _1_An active *cagPAI* was defined by the presence of >17 out of the 27 c*agPAI* genes required for CagA translocation into host cell. _2_*vacA* was classified based on sequence alignments with reference strains G27 (s1b/i1/m1), B8 (s1a/i2/m2) and B38 (s2/i2/m2). None of the Cabo Verde isolates carried the s1a/i1/m1 genotype. hspEuropeCV-C1, hspEuropeCV-Clonal1; hspEuropeCV-C2, hspEuropeCV-Clonal2.

Further clues to the genomic basis of lineage divergence and population expansion of hspEuropeCV-Clonal1 emerged from the analysis of accessory and virulence genes (Figure 3). In contrast to hspEuropeCV-Clonal2, the hspEuropeCV-Clonal1 isolates form a distinct clade in a tree constructed from genetic distances calculated based on accessory gene presence/absence (Figure 3B). Inspection of this tree reveals a cluster of genes that are largely absent in hspEuropeCV-Clonal1 but common in the remaining Cabo Verdean population groups (boxed region in Figure 3C). These genes do not reach significant divergent PBS values due to their intermediate frequency in the parental hspEuropeanCV (Figure 3C). Notably, this cluster corresponds to genes of the cag pathogenicity island (cagPAI), including *cagA*. An inspection of other virulent genes shows that, unlike the other population groups in Cabo Verde, *vacA* in hspEuropeCV-Clonal1 is of the less virulent type s2/i2/m2 (Figure 3C). Finally, considering hosts with single-ancestry colonisations only (n=168), human hosts carrying hspEuropeCV-Clonal1 alone (n=56) tend to have higher pepsinogen ratios than hosts carrying the other lineages (n=112) (Mann-Whitney U = 3865.5, p-value = 0.0103), even after adjusting for host’s age, sex and African ancestry (regression coeff (SE) = 1.77 (0.76), p-value = 0.0213).

Given all of these characteristics of hspEuropeCV-Clonal1, we next carried out a genome-wide association analysis (GWAS) of differentiation between European Cabo Verdean strains with limited evidence of gastric atrophy (pepsinogen ratio > 3) and gastric cancer (GCA) strains (Berthenet et al., 2018) (sample size, geographic origin and bacterial ancestry of Cabo Verde and GCA strains are in Figure 3 – source data 3). In this analysis, we employed the compaction of overlapping k-mers (termed unitigs) approach in DBGWAS (Jaillard et al., 2018). The calculated genomic inflation factor was high (λ=3.4, higher than λ=1-1.1 expected in well-controlled analyses. This inflation is mainly due to differences between hspEuropeCV-Clonal1 and GCA strains: around half of the gene hits present (66% unitigs; 52% of unique genes) showed frequency differences greater than 50% between GCA strains and hspEuropeCV-Clonal1 (Figure 3 – source data 4). Within these hits are well-established *H. pylori* virulence factors such as genes within the cagPAI including *cagA*, genes encoding membrane adhesion proteins (*babA2* and *sabA*) and the vacuolating cytotoxin *vacA*. Overall, there was enrichment for genes involved in lipopolysaccharide and peptidoglycan biosynthesis (COG category M, twice the frequency of the next most represented COG categories P (inorganic ion transport and metabolism) and U (intracellular trafficking, secretion, and vesicular transport); Figure 3 – source data 4).

A notable observation was that 19% (n=37) of the unitigs present in 98% of hspEuropeCV-Clonal1 strains and only in 12% of GCA strains aligned to a non-coding region upstream of a multidrug efflux system ATP-binding protein (C694_RS06300 locus tag in reference sequence 26695 with NCBI reference NC018939.1; (Alfaray et al., 2023)). The DBGWAS authors suggest that this signal may indicate the presence of a putative mobile genetic element in this region (Jaillard et al., 2018). However, searches in ICEfinder (Wang et al., 2024) and ISfinder (Siguier et al., 2006) databases, and prophage analysis with geNomad (Camargo et al., 2024) and PhageBoost (Siren et al., 2021) did not identify any integrative conjugative elements (ICEs), insertion sequences (ISs), or prophage elements in this genomic region. Further investigation will therefore be required to determine whether smaller, or highly diverged mobile genetic elements contribute to the observed signal in this genomic region.

Collectively, these analyses demonstrate that hspEuropeCV-Clonal1 exhibits a distinctive genetic background from GCA strains and provides minimal support for the hypothesis that the expansion of this lineage is driven by increased virulence.

### *H. pylori*-Host codiversification in Cabo Verde

We estimated the individual African ancestry of the human hosts using ADMIXTURE (K=2) with the YRI (Yoruba from Ibadan) and IBR (Iberian from Spain) from the 1000 Genomes Project (Genomes Project et al., 2015) as reference panels. The median human West African ancestry in this cohort is 0.70 (range 0.47-0.83), which is within the range previously reported for the Santiago island, the origin of the majority of the hosts in this sample (based on an independent genomic dataset for 685 Cabo Verdeans across the archipelago, median West African ancestry in Santiago: 0.74, range 0.45-0.86; higher than the median West African ancestry for the total archipelago: 0.58, range 0.24-0.88) (Beleza et al., 2013; Korunes et al., 2022). However, PCA replicates previous patterns of population structure (Figure 4B) (Alexander et al., 2009; Korunes et al., 2022; Laurent et al., 2023) related with host demographic history and island geography.

**Figure 4.**
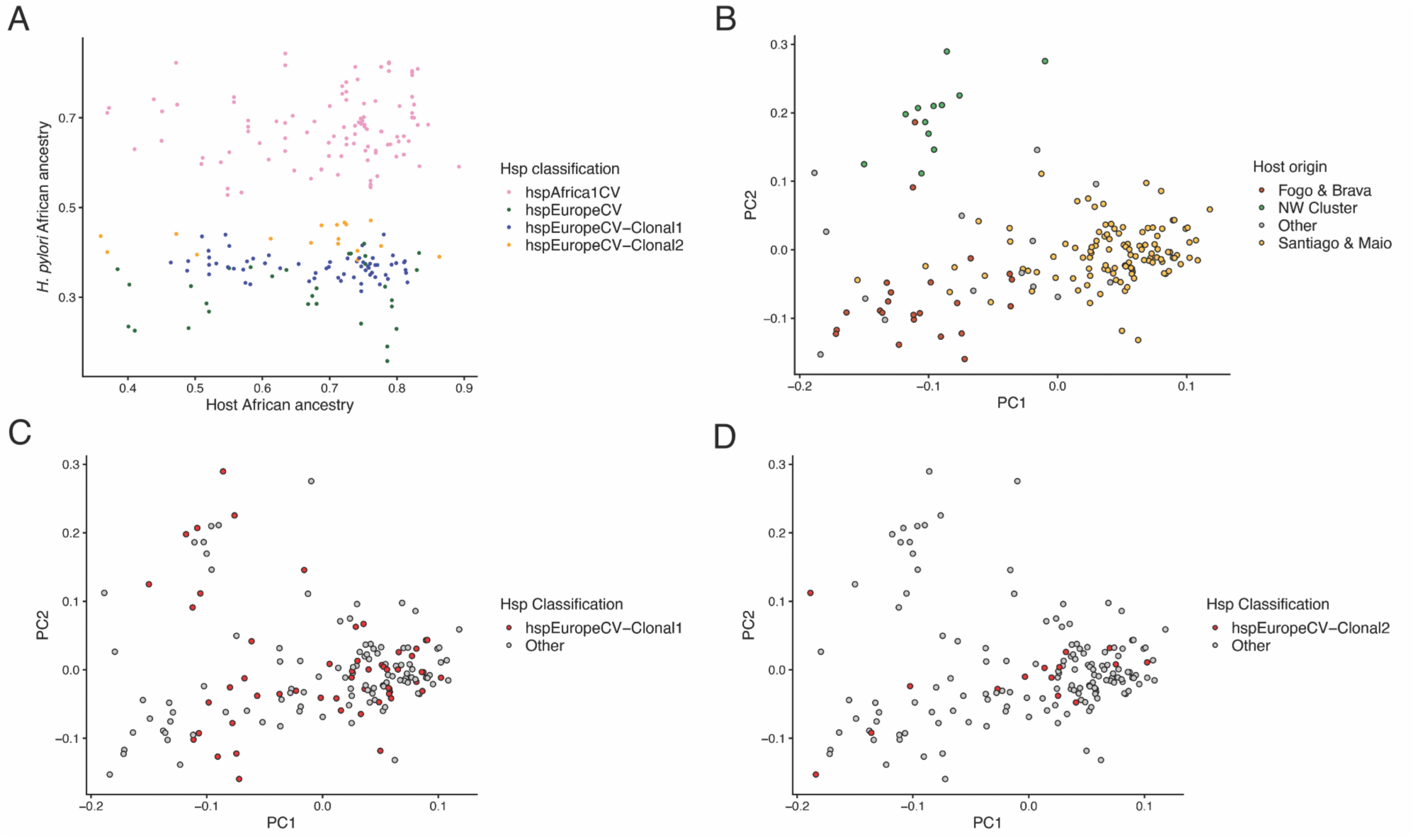
Host-*H. pylori* codiversification in Cabo Verde. (A) Host west African (YRI) ancestry (X-axis) is plotted against *H .pylori* African (hpAfrica1WestAfrica) ancestry (Y-axis). (B) PCA of host genetic variation. Individuals are coloured by parental island group. Other refers to hosts whose parents are from different islands. (C) same PCA as is (B) but with datapoints coloured red to highlight hosts infected with hspEuropeCaboVerde-Clonal1 strains. (D) same PCA as in (B) but with datapoints coloured red to highlight hosts infected with hspEuropeCaboVerde-Clonal2 strains. N_host_ = 174; N*_H.pylori_* = 217.

In this human genomic context, analysis of the relationship between human and infecting *H. pylori* genomic ancestry and structure in Cabo Verde was performed for each *H. pylori* population group. As individual African ancestry in *H. pylori*, we used the estimates obtained with SOURCEFIND because it provides more accurate admixture proportions by correcting for biases introduced by uneven sample sizes among the reference populations (Chacon-Duque et al., 2018).

Notably, the proportion of human West African ancestry was not correlated with the proportion of hspAfrica1WAfrica ancestry overall (n=213, Spearman’s rho= 0.115; p-value= 0.096, Figure 4A), or in any of the *H. pylori* population groups (hspAfrica1CV, n=103, Spearman’s rho= 0.116, p-value = 0.242; hspEuropeCV, n=30, Spearman’s rho= 0.082, p-value = 0.668; hspEuropeCV-Clonal1, n=63, Spearman’s rho= -0.181, p-value = 0.156; hspEuropeCV-Clonal2,n=17, Spearman’s rho= -0.083, p-value = 0.750). Similar results were obtained when the analysis was restricted to hosts carrying a single *H. pylori* strain (total n = 136; n per *H. pylori* population group=[13-51]; all *p-*values > 0.1). Isolates were also distributed across islands with no correlation with host origin or human population group produced in PCA (Figure 4C-D; for each *H. pylori* population group except hspEuropeCaboVerde-Clonal2 for which sample size is too small for statistical hypothesis testing, p-values of Fisher exact tests comparing the number of isolates between northern and southern islands were >0.1). Finally, *H. pylori* population structure was not associated with host’s age, sex, family relationship or household relationship (p-values of Fisher exact tests or logistic regression for >2 categorical data were >0.05). Therefore, there is no evidence for sampling or epidemiological clustering of any of the *H. pylori* population groups.

## Discussion

Our analysis of *H. pylori* in Cabo Verde leverages a unique population sample shaped by founder effects and five centuries of European and African admixture. A major strength of our study lies in the use of samples from hosts from the general population, which offered two primary advantages: firstly, it allowed for an unbiased calculation of seroprevalence and, secondly, it provided a more representative sample of *H. pylori* genetic diversity and host–bacterium combinations within the archipelago to evaluate fully the population structure and evolution of *H. pylori* in the islands.

The estimated *H. pylori* prevalence of 83% (95%CI: 80.0-85.6%) in Cabo Verde (recorded in 2016) is comparable to rates observed in other African nations between 1989 and 2008 (Hooi et al., 2017; Zamani et al., 2018). However, this figure is slightly higher than the mean prevalences reported for several African countries in a large meta-analysis covering the decade 2011–2022 (Li et al., 2023). This discrepancy may be partially explained by observation of an overall decline in the prevalence of *H. pylori* in Africa (Zamani et al., 2018; Li et al., 2023); this trend may not be evident in Cabo Verde due to the absence of longitudinal surveillance data. Additionally, it may also reflect our reliance on a serological method (which aligns with Hoi et al. (2017) and Zamani et al. (2018) but is in contrast with Li et al. (2013)), which does not distinguish active from past infections and tends to lead to higher prevalence estimates (Li et al., 2023). The widespread colonization of people in Cabo Verde, regardless of sex, age and educational levels, highlights a pressing need for further study on risk variants for gastric disease within populations. As we have not evaluated diseased individuals here, future research should investigate whether risk profiles vary among admixed individuals based on their proportion of African ancestry.

Our haplotype- and genotype-based analyses refine our understanding of how the Trans-Atlantic slave trade shaped the subcontinental ancestry of *H. pylori* populations in Cabo Verde and the Americas. By integrating an expanded population reference dataset and restricting African and European source populations to genetically well-resolved lineages, we uncover new patterns of genome variation and admixture. In particular, the identification of a new distinct West–Central African cluster (hspAfrica1WCAfrica) highlights the genetic heterogeneity within African-derived *H. pylori* lineages and provides a parsimonious explanation for the ancestry of many American isolates formerly grouped under hspAfrica1NAmerica (Thorell et al., 2023). In addition, the within-Africa and within-Europe ancestry patterns observed between Cabo Verdean and American strains are consistent with European colonisation and historical migrations during and since the trans-Atlantic slave trade (Ongaro et al., 2019; Micheletti et al., 2020). Together, these findings underscore the need to expand sampling of African *H. pylori* populations and to more comprehensively characterise genetic variation across Africa. Such efforts will allow refinement in the reconstruction of the evolutionary and historical trajectories of *H. pylori* in mixed populations originating during and since the trans-Atlantic slave trade. Importantly, they will also be critical for dissecting patterns of genetic variation underlying disease risk, both within Africa and in African-derived populations.

Genetic variation in the human population of Cabo Verde is predominantly explained by admixture (Korunes et al., 2022). In contrast, ancestry-based population structure is not the main axis of genetic differentiation in the Cabo Verdean *H. pylori*. Instead, the major feature driving population structure in Cabo Verde is the presence of a lineage that most likely arose through a combination of genetic drift and admixture, which then may have subsequently expanded in the population through adaptive evolution. Resolving the relative contributions of genetic drift and selection to observed genetic variation in hspEuropeCV-Clonal1 requires a new statistical framework that evaluates the branching pattern of observed genealogies while accounting for the high recombination rates characteristic of *H. pylori*. Notably, this lineage derives from a European ancestral background and has evolved towards a carriage-associated state (whose ability to induce host gastric atrophy remains to be evaluated) showing significant divergence from gastric cancer strains. The fact that these strains were isolated from individuals of the general population rather than from patients referred to endoscopy, enabled the characterization of gene variants that are likely overlooked in clinical collections enriched for gastric disease.

Frequent cross-infection between unrelated individuals, admixture and clonal expansions have broken the correlation between host and *H. pylori* ancestries in the general population of Cabo Verde. This contrasts with findings from symptomatic individuals in Colombia (Kodaman et al., 2014), where a positive ancestry correlation between *H. pylori* and human host remains. However, to fully contrast with Kodaman et al. (2014), we would need to evaluate if this relationship is similarly disrupted in diseased Cabo Verdean individuals. More broadly, the discrepancy in ancestry correlation between studies highlights the stochastic nature of admixture and its role in generating distinct coevolutionary trajectories that can lead to different phenotypic outcomes. The disrupted host-*H. pylori* ancestry correlation in Cabo Verde indicates that admixture has been extensive enough to decouple the co-occurrence of specific human-*H. pylori* variants that are due to parallel ancestral lineages. As such, Cabo Verde represents a valuable natural system for investigating how host–*H. pylori* genetic interactions influence disease risk and progression.

## Supporting information

Archampong_etal_Suppl_material

## Data availability

All *H. pylori* genomic datasets used here, including the new genomes sequenced in this study, are available under accession numbers provided in Figure1-Source Data 1. The 26695 reference sequence is available on NCBI under Ref Seq accession NC018939.1 and assembly accession CP003904.1.

## Acknowledgments

This work was funded by: the Medical Research Council (MRC-UK), with grant number MR/M01987X/1 awarded to SB; the Biotechnology and Biological Sciences Research Council (BBSRC-UK) and the Midlands Integrative Biosciences Training Partnership (MIBTP), grant numbers BB/M01116X/1 awarded to YLT and CD, and BB/T00746X/1 awarded to IB; and a University of Leicester Doctoral Program Training Fellowship awarded to ST. The funders had no role in study design, data collection and analysis, decision to publish, or preparation of the manuscript.

This research used the ALICE High Performance Computing facility at the University of Leicester. We thank Dr Roxana Zamudio-Zea for her technical assistance with laboratory experiments, Dr Ebony Cave Cave for bioinformatics support, and Isandra Pina for her support at the early stages of sample collection. We are especially grateful to Dr. Dulce Dupret for providing invaluable information regarding gastric pathologies in Cabo Verde.

We also sincerely thank Dr. Agostinho Neto Hospital, Cabo Verde’s Health Department, the Ponta D’Água Health Centre, the Tira-Chapéu Health Centre, and the University of Cabo Verde for providing the facilities and professional support during data collection and follow-up of the participants. Finally, we are deeply grateful to the Cabo Verdean population for their participation in this study.

## Material and Methods

### Ethics

The study was approved by the National Ethical Committee for Health Research of Cabo Verde (Deliberation n°19/2014) and by the Ethics Sub-Committee for Medicine and Biological Sciences of the University of Leicester (protocol 4996-sdsb1-genetics). All participants provided written consent after an explanation of the goal and procedures of the study. In the global population analysis, we additionally incorporated newly generated genome-wide sequences from four Ghanaian isolates obtained from patients presenting upper gastrointestinal symptoms, and 47 genome-wide sequences of Portuguese isolates from patients with gastritis. All isolates were obtained with participants’ informed consent and under ethics approval by the Department of Medicine, Korle-Bu Teaching Hospital, Accra, Ghana (protocol CHS-Et/M.6 - P 4.6/2015-2016), and by the University Hospital Centre (CHU), Porto, Portugal (protocol 003-DEFI/004-CES/2009).

### Sample collection, determination of *H. pylori* serostatus and indirect measurement of mucosal damage

Samples were collected in the capital city Praia, Santiago Island, Cabo Verde from February 2016 to April 2017. A participant was considered of Cabo Verdean origin only if the participant and their parents were native from Cabo Verde. All individuals of both sexes and >20 years of age were invited to participate regardless of showing *H. pylori* infection-related symptoms. Individuals that had self-reported use of antibiotics in the six months prior to sampling were excluded. All individuals provided saliva, blood and gastric mucus samples. Standard questionnaires were used to collect personal and family demographic data (age, sex, place of birth, education level) and *H. pylori* related symptoms (dyspeptic symptoms such as epigastric pain, nausea, and vomiting). An individual’s island origin was defined based on their and their parents place of birth.

*H. pylori* status was determined by testing for IgG against *H. pylori* in the serum of centrifuged samples, using the human anti-*Helicobacter pylori* IgG ELISA Kit (Biohit Healthcare, Finland). The kit reports a sensitivity of 91% and a specificity of 89%. *H. pylori* IgG concentration in serum was also calculated from comparison with a standard curve drawn with control samples included in the ELISA kit.

Indirect measurement of gastric mucosal inflammation and atrophy were assessed by ELISA quantification of the blood biomarkers pepsinogen I and II (PgI and PgII; Gastroview kit, Biohit Healthcare, Finland; (Telaranta-Keerie et al., 2010). The kit defines cut-offs of PgI <30 µg/L <u>and/or</u> PgI/PgII ratio<3 as indicative of atrophic gastritis (Telaranta-Keerie et al., 2010).

The full participant flow is in Figure 1-figure supplement1.

### Epidemiological assessment

We used *H. pylori* serostatus to assess the prevalence of *H. pylori* colonization in Cabo Verde and association with covariates previously described as risk factors, such as age, sex, education level (de Martel and Parsonnet 2006; Wang et al., 2025). Whenever possible (age, education level – see below, pepsinogens-see below), we defined covariates as categorial and continuous variables, and tested associations between variables and *H. pylori* serostatus using logistic and linear regression. The results were consistent between the two regression models. All analyses were performed considering unrelated individuals (hosts) only.

Educational level was evaluated using categorical and continuous variables. The categorical variable considered three levels: lower education (participant did not receive any formal education or received elementary schooling only); intermediate education (participant received secondary schooling); higher education (participant received higher vocational schooling, or university and/or postgraduate education). The continuous variable consisted in the number of years in school education.

To assess gastric inflammation and atrophic gastritis, the levels of PgI, PgII, and their ratio (PgI/PgII) were compared between seropositive and seronegative individuals by regression analysis. We also evaluated how cut-offs for atrophic gastritis defined by the kit were associated with *H. pylori* serostatus using the Kruskal-Wallis test for continuous variables and the Pearson’s chi-square test for categorical variables. Statistical analyses were performed in R.

### *H. pylori* isolation and culture

Gastric mucus samples were collected for *H. pylori* isolation with the minimally invasive “Gastric String Test”, based on the (Velapatiño et al., 2006) protocol. In brief, a 90cm cotton fibre in a gelatine capsule was swallowed by the fasting participant, but with one of the ends of the fibre being taped onto the subject’s cheek. Following one hour, the string was retrieved and the distal 30cm that were in contact with the stomach was placed in transport medium (Brain heart infusion (BHI) broth (Oxoid, UK), 20% glycerol (Thermo Fisher Scientific, UK), 1% Skirrow antibiotic supplement (11.5 μg/ml vancomycin, 5.8 μg/ml Trimethoprim, 0.3 μg/ml Polymyxin B; Oxoid, UK), and 5µg/ml Amphotericin B (Thermo Fisher Scientific), and frozen in dry ice.

During culture, string and transport medium were vortexed vigorously for three minutes and then plated onto two selective BHI blood agar plates containing 5% brain heart infusion agar (BHIA), 7% defibrinated horse blood (Thermo Fisher Scientific), 1% BD BBL IsoVitaleX enrichment media (Thermo Fisher Scientific), 1% Skirrow antibiotic supplement, plus one of two antibiotic mixes: 5 µg/mL Amphotericin B only; or 5 µg/mL Amphotericin B and 30µg/mL Bacitracin. Samples were incubated for up to 10 days at 37°C, in microaerophilic conditions established by GENbag-GENbox systems (bioMérieux France). All small transparent, catalase and urease positive colonies were expanded and confirmed to be *H. pylori* colonies by PCR amplification of a minimum of two and a maximum of three housekeeping genes according to PubMLST amplification primers and conditions (https://pubmlst.org/organisms/helicobacter-pylori/primers). Bacterial DNA was extracted using the phenol/chloroform method (Chen and Kuo 1993), with a modified initial lysis buffer composed by 40 mM Tris-Acetate (pH 9.0), 20 mM sodium acetate (pH 9.0), 60 mM EDTA (pH 9.0,) 1% SDS, to efficiently inactivate *H. pylori* nucleases.

### *H. pylori* genome sequencing, read mapping and SNV calling

*H. pylori* genomic DNA was paired-end sequenced (Illumina HiSeq 4000 at Oxford Genomics Centre) to a mean coverage of 80x. Raw FASTQ reads were trimmed and filtered with Trimmomatic 2.0 using software specifications (Bolger et al., 2014), and quality-checked using FastQC 0.11.5 (Andrews 2010). Paired reads were mapped to the *H. pylori* 26695 reference genome (NCBI Ref Seq accession: NC018939.1; assembly accession: CP003904.1), and SNVs were called using Snippy v4.6.0 (Seemann 2015). SNV annotation was performed with snpEFF (Cingolani et al., 2012), via Snippy by adding the flag ‘--reference’ and providing the reference’s Genebank file. Bcftools v1.2.0 (Danecek et al., 2021) was then used to merge isolates’ VCF files and to subset on biallelic SNVs with the flags **‘**-m2 -M2 -v snps’. To identify core genome SNVs, we ran PLINK v1.90b3 (Chang et al., 2015) with ‘--freq’ to extract the genomic positions of SNVs that were present in either 100% (for population and ancestry analyses) or 95% (for per SNV and per gene F_ST_ and PBS analyses) of chromosomes. The resulting list of core positions was then used to subset the VCF file with VCFtools v1.14.0 (Danecek et al., 2011).

### H. pylori de novo assembly

Trimmed, paired reads were assembled to contig level using SPAdes 3.12 with both mismatch repair and coverage threshold (set to auto) parameters implemented (Bankevich et al., 2012). Contigs were further assembled into scaffolds using the “assembly improvement pipeline” from Sanger Pathogens (Page, De Silva, et al., 2016). Assembly quality was assessed via QUAST 4.3.

The resulting draft genomes were annotated and aligned using Prokka v1.14.6 (Seemann 2014) and PIRATE v1.0.5 (Bayliss et al., 2019), considering 50, 70, 90 and 95% sequencing identity. Subsequently, core genomes were built as a concatenation of genes identified to be present in 100% of input sequences using PIRATE’s output files and python script ‘create_pangenome_alignment.pl’ (found in PIRATE Github’s page https://github.com/SionBayliss/PIRATE/tree/master/scripts), with the parameters -- dosage 1.25 -t 100 -I PIRATE.gene_families.ordered.tsv -f ./feature_sequences/.

### *H. pylori* reference dataset and genome alignment

Complete genomes of 1705 global *H. pylori* isolates were retrieved from public repositories (NCBI, Datadryad; last accessed December 2024) to investigate worldwide population structure and genetic ancestry of *H. pylori* populations (Figure 1 – source data 1). This comprehensive dataset comprises isolates whose population of affiliation has been assessed previously and have been classified into major populations (“hp”) and subpopulations (“hsp”), including those from the *Helicobacter pylori* Genome Project (HpGP; (Thorell et al., 2023)), as well as isolates whose population of affiliation has not been previously assessed, four new Ghanaian genomes and 47 new Portuguese genomes from our own collection.

PIRATE genome alignments were quality-checked using QUAST v4.3 (Gurevich et al., 2013), and only assemblies with a N50 below 20kb were excluded; this single quality control (QC) threshold enabled the inclusion of a broader representation of certain South America countries, as well as all African genomes available up to December 2024. Sequence annotation and alignment of these sequences together with the Cabo Verde dataset was done using Prokka v1.14.6 and PIRATE v1.0.5 as described above. SNP calling for each alignment was performed using SNP-sites (Page, Taylor, et al., 2016).

A preliminary analysis of Single Nucleotide Variant (SNV) differences between pairs of aligned genomes with SNP-dists (Seemann 2021) showed the presence of clonal or closely related isolate pairs (characterized by an excess of isolates falling within the first percentile of the pairwise SNV difference distribution (<1000 pairwise SNV differences) and mostly representing duplicated samples in the database, within-host comparisons, or related individual comparisons). To enable a more robust analysis of deeper evolutionary relationships among *H. pylori* strains, we have cleared the dataset of these recent, clonal, relationships. Figure 1 – source data 1 includes only those isolates that passed QUAST QC, were determined to be unrelated (not clonal) and that have less than 2% missing genotypes.

### Population genetics and ancestry analyses in a global context

We first defined a reference dataset comprising isolates known to belong to hpAfrica1, hpAfrica2, hpEurope, hpAsia2, hspEAsia, hspAmerind and confirmed to have originated from corresponding geographic regions. We then carried out k-means clustering with the KMeans algorithm in Python’s scikit-learn package (Pedregosa et al., 2012)(scripts can be found in (Cave 2025)) and rounds of PCA using PLINK v1.9 to further resolve the continental and subcontinental ancestry of unclassified isolates (Figure 1 – figure supplement 3). For more computationally efficient population genetic analyses of African, European, American and Cabo Verde isolates, we subsetted the European dataset to a final total sample of 459 isolates. The resulting alignment length was 860,079 bp.

We used fineSTRUCUTRE v4 to run the complete ChromoPainter/fineSTRUCUTRE analysis pipeline (Lawson et al., 2012). FineSTRUCTURE v4 was run with default parameters: 50,000 iterations of both the burn-in and Markov chain Monte Carlo (MCMC) model. In preliminary analyses, we included genetic variants with ≤ 2% missing genotype data, and carried out genetic imputation using SHAPEIT v2.r837 (Delaneau et al., 2008) with parameters --states 500 --burn 10 --main 50 --prune 10. The ‘makeuniformrecfile.pl’ (part of the fineSTRUCTURE v4.1.1 package) was used to produce a uniform recombination map across *H. pylori* genomes. To analyse population structure in detail, we ran PCA on the coancestry matrix from the “chunkcounts.out” output file. The matrix was first normalised by setting the diagonal to the average of each row, then zero meaning by subtracting the row means. Subsequent fineSTRUCTURE analyses (excluding outlying isolates in Figure 1 -figure supplement 3) were performed using only genetic variants with complete genotype data. The final fineSTRUCTURE analysis contained 1165 isolates and 37,480 SNVs (Figure 1 – source data 1).

ADMXITURE v1.3.0 (Alexander et al., 2009) was ran for values of K=2-9 in the same final dataset. We pruned the data to remove sites in high linkage disequilibrium (LD) using PLINK v1.90b3 with parameters --indep-pairwise 50 10 0.1. A total of 16,910 SNVs with no missing genotypes was included in the analysis.

### Population genetics and ancestry analyses in Cabo Verde

A phylogenetic tree was built from PIRATE alignments using FastTree v2.1 (Price et al., 2010) with the generalised time-reversible model of nucleotide evolution and the discrete gamma model with 20 rate categories for rescaling. The tree was visualised in iTOL v5 (Letunic and Bork 2021). PCA was performed in PLINK v1.9.

For ancestry analyses, we ran fineSTRUCTURE V4.1.1 with PIRATE alignments incorporating hspAfrica1WAfrica and hspSWEurope as ancestral population references. This analysis included a total of 212,074 SNV with no missing genotypes. The resulting coancestry matrix from the “chunkcounts.out” output file was visualised as a heat map, using the authors’ R functions available at https://people.maths.bris.ac.uk/∼madjl/finestructure/finestructureR.html.

To calculate population and individual level ancestry in Cabo Verdean *H. pylori* while accounting for differences in the reference sample sizes, we ran SOURCEFIND v2 (Chacon-Duque et al., 2018) with the ‘*.chunklengths.out’ output file from fineSTRUCTURE and using hspSWEurope and hspAfrica1WAfrica as surrogates. As recipients, we either included the two reference populations and the four Cabo Verdean population groups (for population-level ancestry estimates), or each Cabo Verdean isolate individually (for individual-level ancestry estimates). In all SOURCEFIND runs, we fixed the truncated Poisson prior to four surrogate groups per individual, allowing up to eight groups to contribute ancestry at each MCMC iteration. We performed 2,000,000 iterations with a 50,000-step burn-in, sampling mixture coefficients every 5,000 iterations. Final ancestry proportions were calculated as the mean of all posterior samples.

Hierarchical clustering based on the accessory genomes’ presence /absence of genes was performed with the pheatmap v1.0.13 package in R v4.5.2, using the complete linkage method. Only genes present between 95% and 5% of the isolates (n=414), as determined from the PIRATE pangenome alignment, were used in this analysis.

### ABC inference

We modelled the demographic history of three of the four Cabo Verdean population groups with adequate sample size using the ABC framework from (Davison et al., 2024), with the following modifications. For each population group, we generated a reference table of 100,000 simulated SNP datasets with the same sample size and number of synonymous (neutral) SNVs, *S*, as the observed genome-wide (hspAfrica1CV, *S* = 138,091; hspEuropeCV, *S* = 153,087; hspEuropeCV-Clonal1, S = 143,501), and whose SFS is derived from an analytical approach (Freund et al., 2023) that incorporates the Kingman coalescence model with exponential growth characterized by a specified growth rate *g*. Specified growth rates were drawn from uniform prior distributions on the discrete sets of [0-15] for hspAfrica1CV and hspEuropeCV, and [0-30] for hspEuropeCV-Clonal1, increasing by a step size of 0.05 (note that these included no growth (*g* = 0), that is constant population sized). For all simulated and observed datasets, we summarised genetic diversity using the 10% percentiles of pairwise Hamming distances between sequences, scaled by their maximum within the sample (simulated or observed), and Tajima’s D. Calculations on observed data were based on synonymous (neutral) variation. We then compared the summary statistics between simulated and observed dataset using the Random Forest ABC classifier (ABC-RF) from the *abcrf* R-package (Raynal et al., 2019), which provided the posterior distributions of the populations’ growth-rate parameter. The summary statistics with higher discriminatory power was Tajima’s D. Notably, with this approach, the posterior estimates of the growth rate are given in *Ne* coalescent time units (*Ne*, population effective size), which means that growth happens on large timescales (especifically by exp(g/*Ne* * growth rate estimate)). To evaluate the accuracy of the model the normalized mean absolute error (NMAE), which averages |estimated value–true value|/true value across all simulations, for all other true growth rates combined.

To assess whether the model fits the observed genetic diversity, we performed graphical posterior predictive checks by calculating the Tajima’s D of 1,500 simulated genomes, obtained with growth rates drawn uniformly from the full-range ABC-RF posterior quantiles at intervals of 0.025 (Figure 2-Figure supplement 3).

### Per SNP and per gene FST, PBS, and COG-function enrichment analysis

We calculated pairwise F_ST_ values for each SNV within the core genome and for the presence/absence profiles of accessory genes. We utilized Hudson’s F_ST_ estimator, as it is less sensitive to differences in sample size and SNV ascertainment schemes (Bhatia et al., 2013). The core SNV F_ST_ values were computed using the scikit-allel Python package (Alistair and Harding 2017), while the F_ST_ values for accessory genes were calculated using a custom implementation of the Hudson formula in R. Core SNVs were called via mapping to the reference sequence 26695 and correspond to those present in ≥95% of the total sample. Accessory genes were obtained through PIRATE’s pangenome alignment and correspond to genes present in 5-95% of the total sample.

We then used these F_ST_s to calculate Population Branch Statistic (PBS) values as in Yi et al. (2010). PBS summarizes a three-way comparison of F_ST_s between a focal group (hspEuropeCV-Clonal1 and hspEuropeCV-Clonal2), a closely related population (hspEuropeCV) and an outgroup (hspAfrica1CV). Within the core genome, we focused our analysis on the mean PBS of non-synonymous SNVs, as these variants are more likely to hold functional significance. A core gene was identified as significantly differentiated in the hspEuropeCV-Clonal1 and hspEuropeCV-Clonal2 lineages if it met two criteria: first, that the proportion of its non-synonymous SNVs falling within the top 5% of the PBS distribution exceeded 15% and 10%, respectively (95th percentile of proportions); and second, that its overall mean non-synonymous PBS value ranked within the top 5% of all mean non-synonymous PBS values (Figure 3 – source data 3-4). An accessory gene was identified as significantly differentiated in the hspEuropeCV-Clonal1 and hspEuropeCV-Clonal2 lineages if its PBS values ranked within the top 5% of all PBS values.

Next, we used Eggnog v5.0 (Huerta-Cepas et al., 2019), to identify orthologous groups for each gene and assign them to specific Cluster of Orthologous groups (COG) functional categories. This classification enabled a fold enrichment analysis using Fisher’s exact test to identify overrepresented COG functional categories within the sets of most significantly differentiated core and accessory genes.

### GWAS of genetic differentiation with gastric cancer strains

We collected 53 gastric cancer strains from Bethernet et al. (2018; Figure 3 – source data 4) and used these as cases in a GWAS. Cabo Verdean European strains isolated from hosts with pepsinogen ratios >3 were used as controls. Host mean age was 42.1 years, age range 23 – 86 years, with 75.1 % female.

Associations between genetic variants in both the core and accessory genomes and the binary case–control trait were tested using a De Bruijn graph–based GWAS (DBGWAS) approach implemented in DBGWAS v0.5.3 (Jaillard et al., 2018). DBGWAS is a k-mer– based method that represents sequence variation using compact De Bruijn graphs (cDBGs), in which nodes correspond to DNA fragments of variable length generated by the compaction of overlapping k-mers (termed unitigs). This graph-based representation substantially reduces k-mer redundancy and captures genomic diversity of k-mers within the broader genomic context of the study population. DBGWAS tests for both locus-specific (unitig) and lineage effects on the phenotype using a linear mixed model (LMM), thereby accounting for population structure. This approach controls for population structure by incorporating a genetic relatedness (kinship) matrix in a linear mixed model. DBGWAS was run with default settings and the “-SFF 200” flag which generates the first 200 most significant unitigs. DBGWAS outputs Wald statistics that represent signed Z-scores (Z). To estimate the genomic inflation factor (the λ factor), we calculated median(Z^2^)/0.45494.

All unitig hits were analysed using BLAST v2.13.0 (Altschul et al., 1990) to identify corresponding genes. Eggnog v5.0 (Huerta-Cepas et al., 2019) was subsequently used to assign each gene to orthologous groups and to classify them into COG functional categories.

We also ran the hspEuropeCV-Clonal1 control isolates’ improved *de novo* assemblies through the ICEfinder v2.0 (Wang et al., 2024) and ISfinder (Siguier et al., 2006) databases to identify ICEs and ISs that could be found in the non-coding region upstream of a multidrug efflux system ATP-binding protein (C694RS06300 locus tag in reference sequence 26695 with NCBI reference NC018939.1).

To identify prophages in the same genomic region we followed the pipeline from (Cave 2025). Briefly, the machine learning algorithms of geNomad v1.7.5 (Camargo et al., 2024) and PhageBoost v0.1.7 (Siren et al., 2021) were employed, using Prokka-annotated fasta files as inputs. CheckV v1.5 (Nayfach et al., 2021) was subsequently applied to assess the quality of the identified putative prophage elements. Only elements longer than 20,000 bp, containing at least one viral gene, and with a CheckV-estimated completeness greater than 50% were retained as prophages.

### Human DNA collection, extraction, and sequencing

Human DNA was obtained from saliva and blood samples. Saliva was collected with Oragene DNA 600 saliva kits (DNA Genotek, Canada). Saliva DNA was extracted from 0.5ml of sample with the PrepIT-L2P|PT-L2P reagent according to the manufacturer’s protocol (DNA Genotek) and minor changes: RNA was degraded via introduction of a 30 minute, 37°C incubation with 10mg/ml RNAseA (Thermo Fisher Scientific) after initial DNA release incubation; ethanol volume was increased to 1.5X the supernatant volume for DNA precipitation; ethanol wash strength was increased to 85%; and final resuspension was done in 10mM Tris HCl pH8.5. Blood samples were collected by venepuncture using a Vacutainer® Plastic K2EDTA tube with Lavender BD Hemogard™ Closure 6ml 13×100mm and Safety-Lok™ Blood Collection Set with Pre-attached Holder 21G 0.75” 12” tubing (Becton Dickinson, UK). DNA was extracted via the phenol-chloroform method.

Human DNA samples were whole genome sequenced to 1x mean coverage (Illumina HiSeq 4000) and imputed by Gencove (New York, USA), using their imputation algorithms for low-pass sequencing to call variants with average imputation r^2^> 0.9.

### Human population genetics and ancestry analysis

Biallelic VCFs were filtered based on linkage disequilibrium, missingness per marker, Hardy-Weinberg Equilibrium and minor allele frequency using PLINK 1.9 (PLINK flags:--indep-pairwise 50 5 0.5, --geno 0.1, --hwe 1e-6, --maf 0.1). Remaining variants (660,244 SNVs) were used to calculate PCA using PLINK 1.9 and host ancestry using FastNGS admix (Jorsboe et al., 2017) with Yoruban (YRB) and Iberian (IBR) reference samples from the 1000 Genomes Project (Genomes Project et al., 2015). REAP was used to estimate the kinship coefficient between hosts (Thornton et al., 2012).

## Figure supplements

**Table 1 - figure supplement 1**

**Distribution of Anti-*H. pylori* Immunoglobin G (IgG), Pepsinogen I (PGI), and Pepsinogen II (PGII) levels and PGI/PGII ratios in the general population of Cabo Verde.** (A) Distribution of anti–*H. pylori* IgG levels across age, with subjects grouped into 10-year age bins. (B) Distribution of pepsinogen I stratified by serostatus. PgI levels are significantly higher in seropositive individuals (Regression Coefficient, coeff (S.E.) = 18.0 (4.3) µg/L, p-value= 2.5×10^-5^). (C) Distribution of pepsinogen II stratified by serostatus. PgII levels are significantly higher in seropositive individuals (coeff (S.E.) = 3.8 (0.6) µg/L, p-value=5.2×10^-11^). (D) Distribution the ratio pepsinogen I/pepsinogen II (PgI/PgII) stratified by serostatus (coeff (S.E.) = -3.7 (0.64) units, p-value= 1.1×10^-8^; all statistical analyses were controlled for sex and age).

**Figure 1 - Source Data 1** (in .xlsx format)

List of strains used in the evaluation of population structure of worldwide *H. pylori* strains. List of publications at the end of the table. HpGP, *Helicobacter pylori* Genome Project.

**Figure 1- supplement figure 1**

**Participant flow.**

**Figure 1- supplement figure 2**

**Population structure of worldwide *H. pylori* populations, including Cabo Verde.**

Coancestry PCA obtained including the Cabo Verdean dataset and the dataset from Muñoz-Ramirez et al. (2021). Updated HpGP ‘hsp’ nomenclature is indicated on the plot. Data points are coloured according to the results of a K-means clustering analysis (K = 16) based on the first two principal components (PC1 and PC2). The colour key, shown on the right, indicates the correspondence between K-means clusters and the ‘hsp’ nomenclature defined by Muñoz-Ramirez et al. (2021). The K-means analysis partitions European and African genomes into clusters that do not fully correspond to established hsp groupings. Clusters labelled “Unknown” include the majority of American genomes, reflecting their complex European and African ancestry. Additionally, the “Unknown Africa” clusters include several South African genomes scattered among other African clusters, consistent with the complex admixture and population structure characteristic of South African populations (Patterson et al., 2010). The fineSTRUCTURE analysis replicates the patterns reported by Muñoz-Ramirez et al. (2021) and positions Cabo Verdean genomes between European and African genomes.

**Figure 1- supplement figure 3**

**Within-Africa and within-Europe population structure.** (A–B) PCA of African isolates before (A) and after (B) excluding *hspAfrica2*, which represents an outlier relative to Cabo Verdean ancestry. Data points are colour-coded according to their HpGP (‘hsp’) classification. The “Unknown” category includes unclassified strains from The Gambia and Ghana within the hspAfrica1WAfrica cluster; strains from Cameroon, Nigeria, Sudan, and South Africa within the hspAfrica1MiscellaneousAmerica and hspAfrica1SouthAfrica cluster; and South African strains within the hspAfrica2 cluster. (C–D) PCA of European isolates before (C) and after (D) exclusion of outlier populations and delineation of population groups by K-means clustering (K = 3) based on the first two principal components.

**Figure 1- supplement figure 4**

**Population structure and ancestry of *H. pylori* strains from Europe, African, the Americas and Cabo Verde.** Coancestry PCA of 1342 *H. pylori* strains from Europe, Africa, multiple American populations and Cabo Verde. FineSTRUCTURE was ran with a total number of 268,516 SNVs with ≤2% missing genotype data.

**Figure 1- supplement figure 5**

**Population structure and ancestry of *H. pylori* strains from Europe, African, the Americas and Cabo Verde.** fineSTRUCTURE-inferred dendrogram displaying the MCMC-derived tree assignments from the “_tree.xml” file. Highlighted isolates belong to African and European reference isolates: hspAfrica1WAfrica, hspAfrica1WCAfrica, hspAfrica1SAfrica, hspNEurope, hspEurasia, and hspSWEurope. Colour code is defined in Figure1.

**Figure 1- supplement figure 6**

**Population structure and ancestry of *H. pylori* strains from Europe, African, the Americas and Cabo Verde** A) Coancestry PCA of 1161 *H. pylori* strains from Europe, Africa, multiple American populations and Cabo Verde (without the four new genomes from Ghana). B) fineSTRUCTURE-inferred dendrogram displaying the MCMC-derived tree assignments from the “_tree.xml” file obtained from the same analysis as in A). Colour code is A) and in Figure 1.

**Figure 1- supplement figure 7**

**Population structure and ancestry of *H. pylori* strains from Europe, African, the Americas and Cabo Verde.** A) Coancestry PCA of 1130 *H. pylori* strains from Europe, Africa, multiple American populations and Cabo Verde. hspAfrica1WAfrica and hspAfrica1SAfrica were downsampled to 10 samples. B) fineSTRUCTURE-inferred dendrogram displaying the MCMC-derived tree assignments from the “_tree.xml” file obtained from the same analysis as in A). Colour code is A) and in Figure 1.

**Figure 1- supplement figure 8**

**Genome-wide ancestry of *H. pylori* strains involved in the Trans-Atlantic Slave Trade.** Unsupervised ADMIXTURE Clustering at K=2-7. Only European and African population groups with *n* > 10 are included (Figure 1 – source data 1). Labels are as in Figure 1.

**Figure 1- supplement figure 9**

**Population structure and ancestry of *H. pylori* strains involved in the Trans-Atlantic Slave Trade.** Coancestry PCA analysis of 1165 *H. pylor*i strains from Europe, Africa, multiple American populations and Cabo Verde (as in Figure1), with each plot showing only one of the American samples.

**Figure 2 – figure supplement 1**

**Population Structure of *H. pylori* strain in Cabo Verde.** PCA of Cabo Verdean, hspSWEurope and hspAfrica1WAfrica strains.

**Figure 2 – figure supplement 2**

**Genome-wide ancestry of *H. pylori* strains in Cabo Verde.** Individual ancestry calculated with SOURCEFIND v2 and using hspSWEurope and hspAfrica1WAfrica as surrogates. To calculate individual ancestries, we ran SOURCEFIND with one Cabo Verdean isolate as recipient at a time. Blue, hspSWEurope ancestry; Orange, hspAfrica1WAfrica ancestry.

**Figure 2 - source data 1**

**Mean ancestry in Cabo Verdean, European reference and African reference populations.** Mean popuation ancestry was calculated with SOURCEFIND v2. HspSWEurope and hspAfrica1WAfrica were included as surrogates and all recipients (rows) were allowed to copy from all surrogates (columns), including itself (self-copy; highlighted in yellow).

**Figure 2 – figure supplement 3**

**Genetic diversity of *H. pylori* in Cabo Verde.** Pairwise SNV differences distribution in the four population groups in Cabo Verde.

**Figure 2 – figure supplement 4**

**Posterior Predictive checks (PPCs) of the ABC approach implemented to investigate the demographic history of *H. pylori* population groups in Cabo Verde.** Observed genetic diversity was compared with that obtained from 100,000 simulations via a Random Forest ABC classifier (ABC-RF). The NMAEs obtained when inferring the growth model and corresponding growth rates were ≤0.03. We then performed PPCs. Histograms correspond to Tajima’s D values obtained from simulating 1,500 genomes using as parameters with growth rates drawn uniformly from the ABC-RF posterior quantiles at intervals of 0.025 and spanning the full range from 0 to 1. The red vertical dashed line indicates the observed Tajima’s D value. The final (lower-right) panel shows the PPC from an ABC run using a randomly downsampled hspAfrica1CV sample, matched in size to hspEuropeCV-Clonal1. Although the inferred growth rate is slightly higher and the confidence interval is wider than that estimated using the full sample (growth rate estimate = 0.5, 95% CI [0.25, 1.15] vs. 0.2, 95% CI [0.1, 0.5] for the full sample), the inferred model remains the same.

**Figure 3 – source data1** (in .xlsx format)

**List of differentiated genes in hspEuropeCV-Clonal1.**

**Figure 3 – source data2** (in .xlsx format)

**List of differentiated genes in hspEuropeCV-Clonal2.**

**Figure 3 – source data 4** (in .xlsx format)

**List of strains used in the DBGWAS for a GWAS for Gastric Cancer, using the Cabo Verdean European isolates as controls.**

**Figure 3 – source data 5** (in .xlsx format)

**Top 200 genes from a GWAS for Gastric Cancer, using the Cabo Verdean European isolates as controls.** The table presents, for each unitig, the unitig absolute frequency; unitig’s absolute and relative frequencies in controls, in hspEurope-CV alone, and in cases; the q-value (the Benjamini-Hochberg derived p-value controlling for the false discovery rate); the estimated effect size (EstEff); the Wald Statistic (WaldStat; the significance of the fixed effect of a variant while accounting for population structure), the unitg sequence and length; the gene it aligns to; the gene’s predictive function; the gene’s associated COG category, as inferred by eggnog V5; and variant type (Type; LPG: Local polymorphism in gene; LPN: Local polymorphism in non-coding region of gene). Highlighted in green are genes in common with Bethernet et al., 2018.

