## Supplementary material for "Genomic variation and ancestry of *Helicobacter pylori* in the admixed population of Cabo Verde reveal host adaptation, limited host–pathogen ancestry concordance, and signatures of trans-Atlantic slave trade migrations": Archampong_etal_Suppl_material: elife_Figure 2 - source data 1_revised.docx

**Archampong et al. 2026.**

**Figure 2 - source data 1**

**Mean ancestry in Cabo Verdean, European reference and African reference populations**. Mean population ancestry was calculated with SOURCEFIND v2. HspSWEurope and hspAfrica1WAfrica were included as surrogates and all recipients (rows) were allowed to copy from all surrogates (columns), including itself (self-copy; diagonal, highlighted in yellow).

| **hsp** | **hspAfrica1WAfrica** | **hspSWEurope** | **hspAfrica1CV** | **hspEuropeCV** | **hspEuropeCV-Clonal1** | **hspEuropeCV-Clonal2** |
| --- | --- | --- | --- | --- | --- | --- |
| **hspAfrica1WAfrica** | 0.2032 | 0.185 | 0.2339 | 0.1353 | 0.1171 | 0.1254 |
| **hspSWEurope** | 0.1709 | 0.2626 | 0.1325 | 0.1765 | 0.1292 | 0.1283 |
| **hspAfrica1CV** | 0.2011 | 0.118 | 0.3103 | 0.1295 | 0.1036 | 0.1375 |
| **hspEuropeCV** | 0.1392 | 0.2342 | 0.1279 | 0.2331 | 0.1316 | 0.1339 |
| **hspEuropeCV-Clonal1** | 0.1247 | 0.1378 | 0.1211 | 0.1314 | 0.3877 | 0.0972 |
| **hspEuropeCV-Clonal2** | 0.1363 | 0.1512 | 0.1627 | 0.1487 | 0.1108 | 0.2904 |
