## Supplementary figures and images for "Genomic variation and ancestry of *Helicobacter pylori* in the admixed population of Cabo Verde reveal host adaptation, limited host–pathogen ancestry concordance, and signatures of trans-Atlantic slave trade migrations"

### elife_Figure1-figure_suppl1_revised.pdf

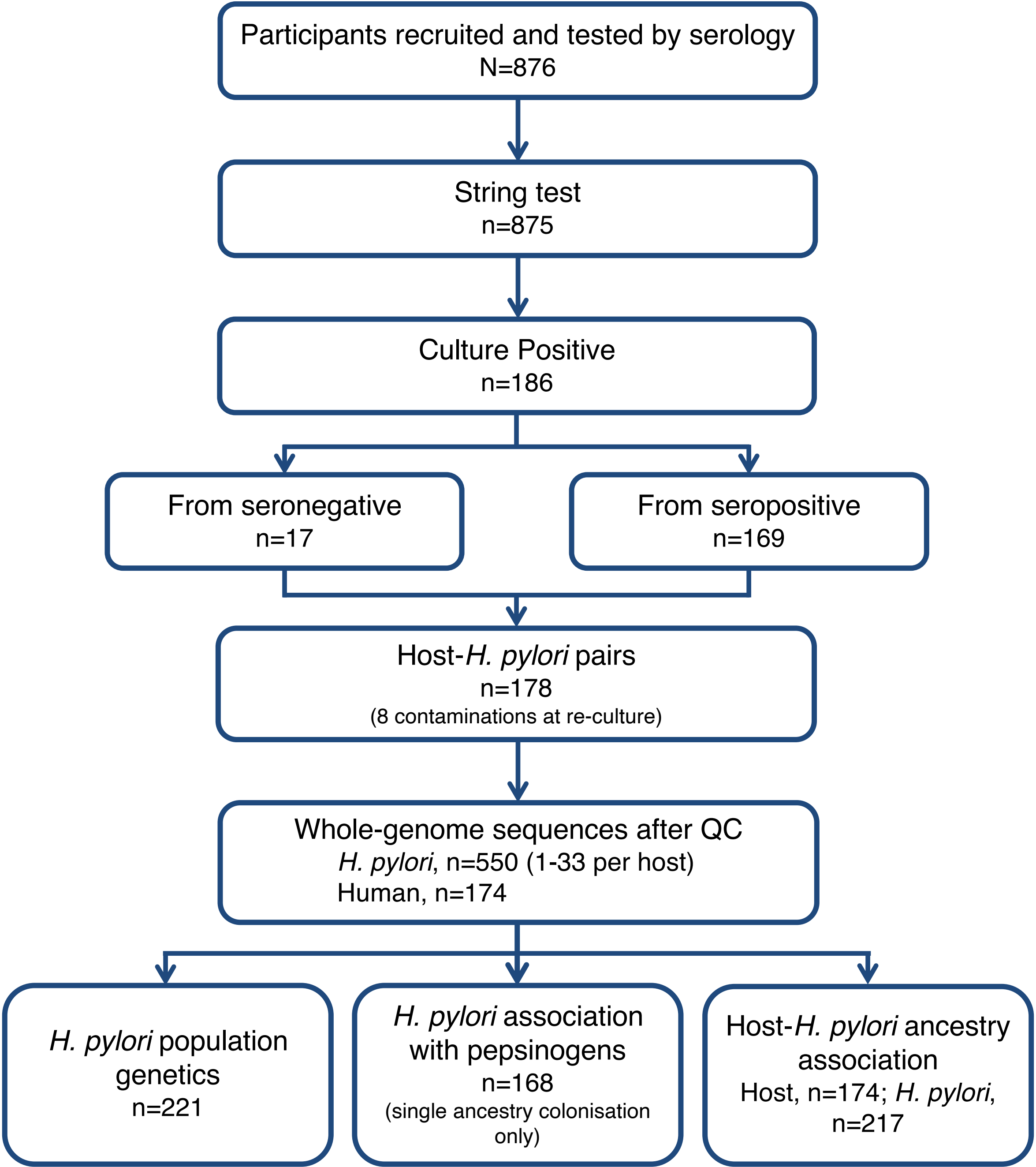

### elife_Figure1-figure_suppl2_revised.pdf

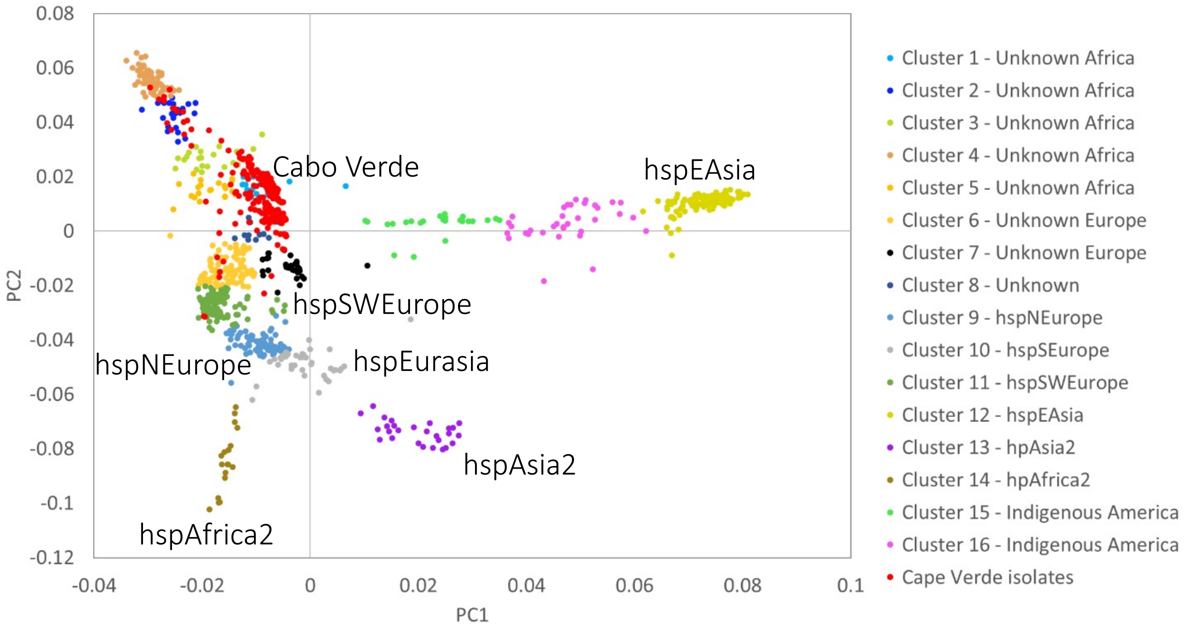

### elife_Figure1-figure_suppl3_revised.pdf

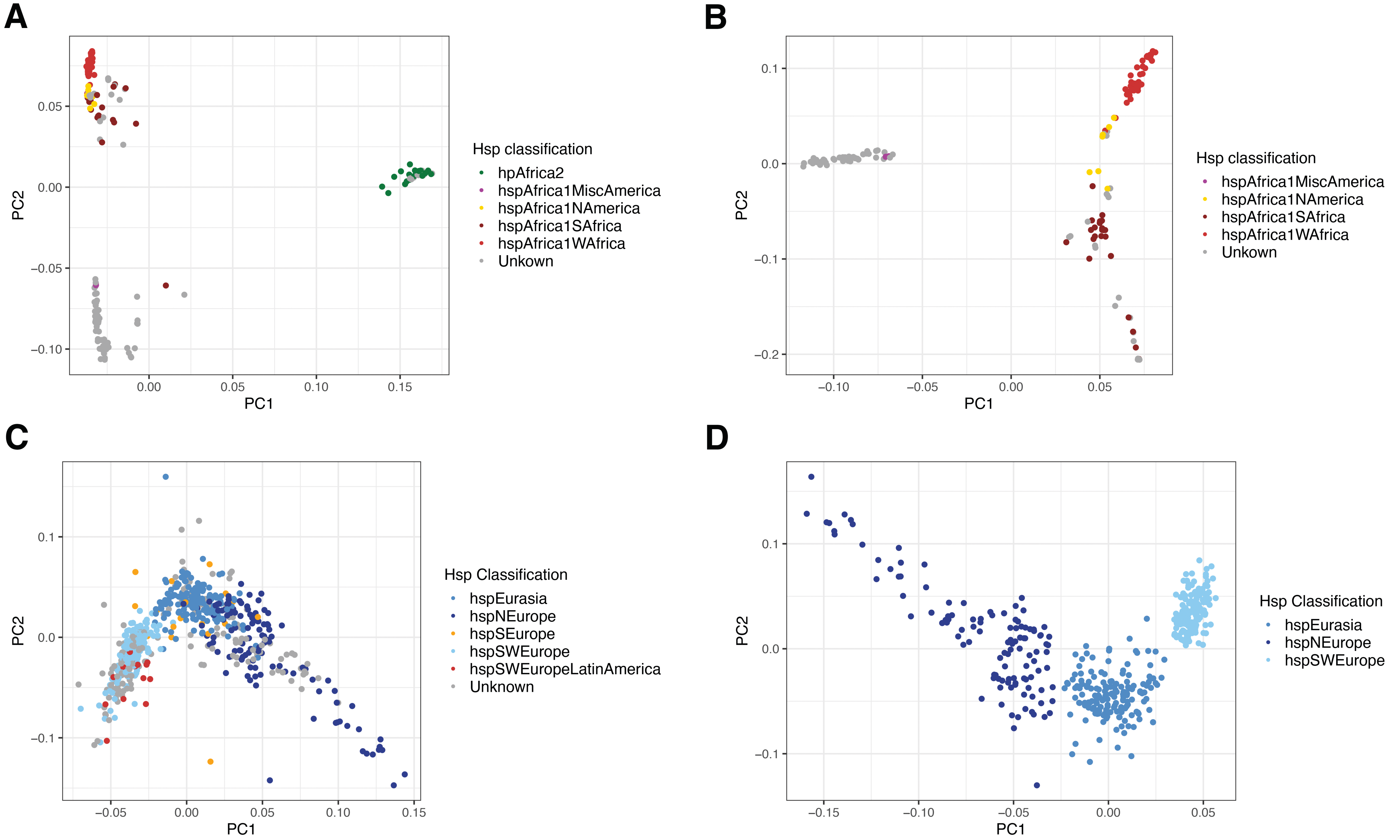

### elife_Figure1-figure_suppl4_revised.pdf

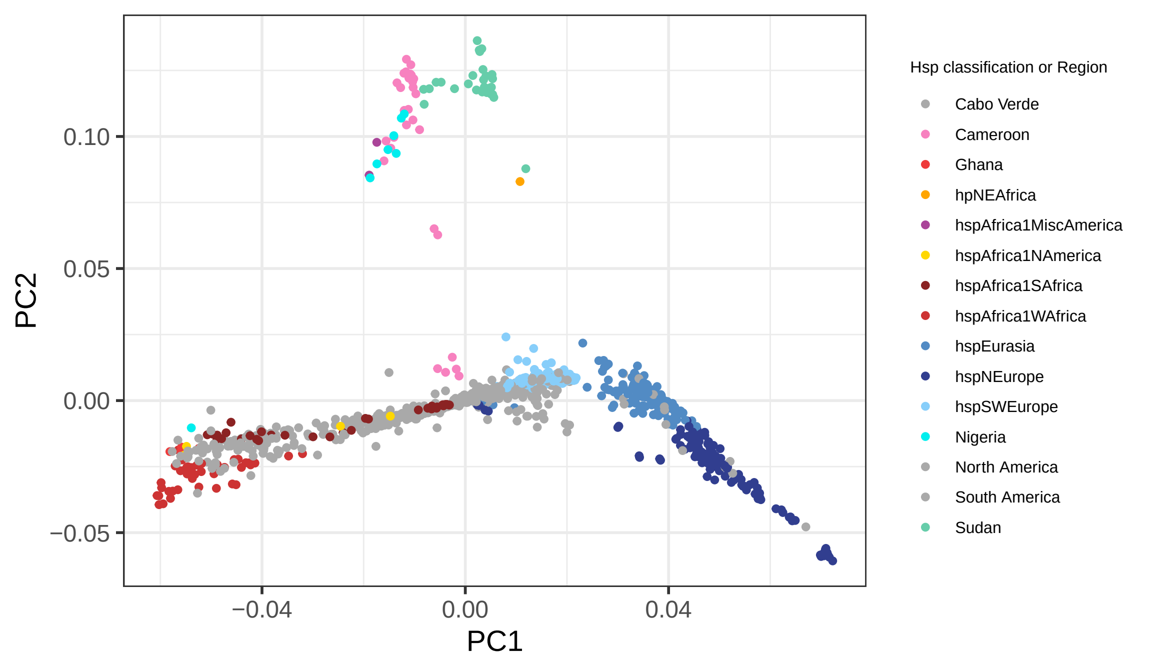

### elife_Figure1-figure_suppl5_revised.pdf

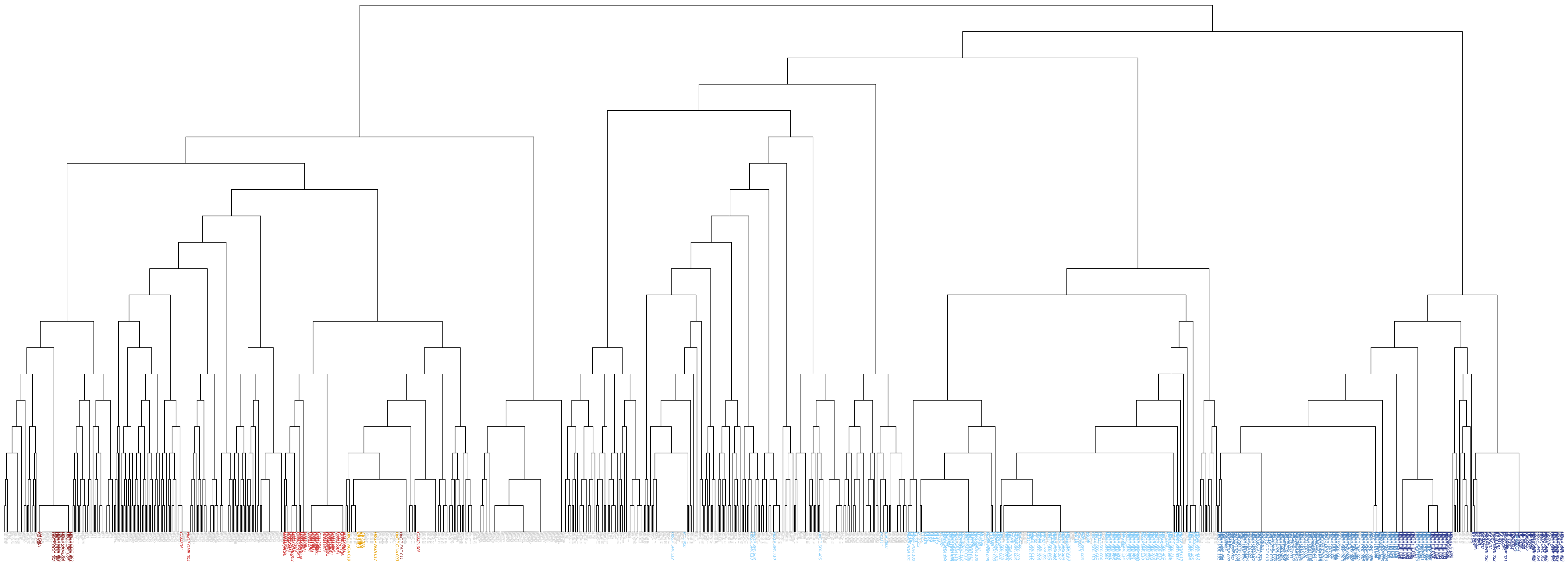

### elife_Figure1-figure_suppl6_revised.pdf

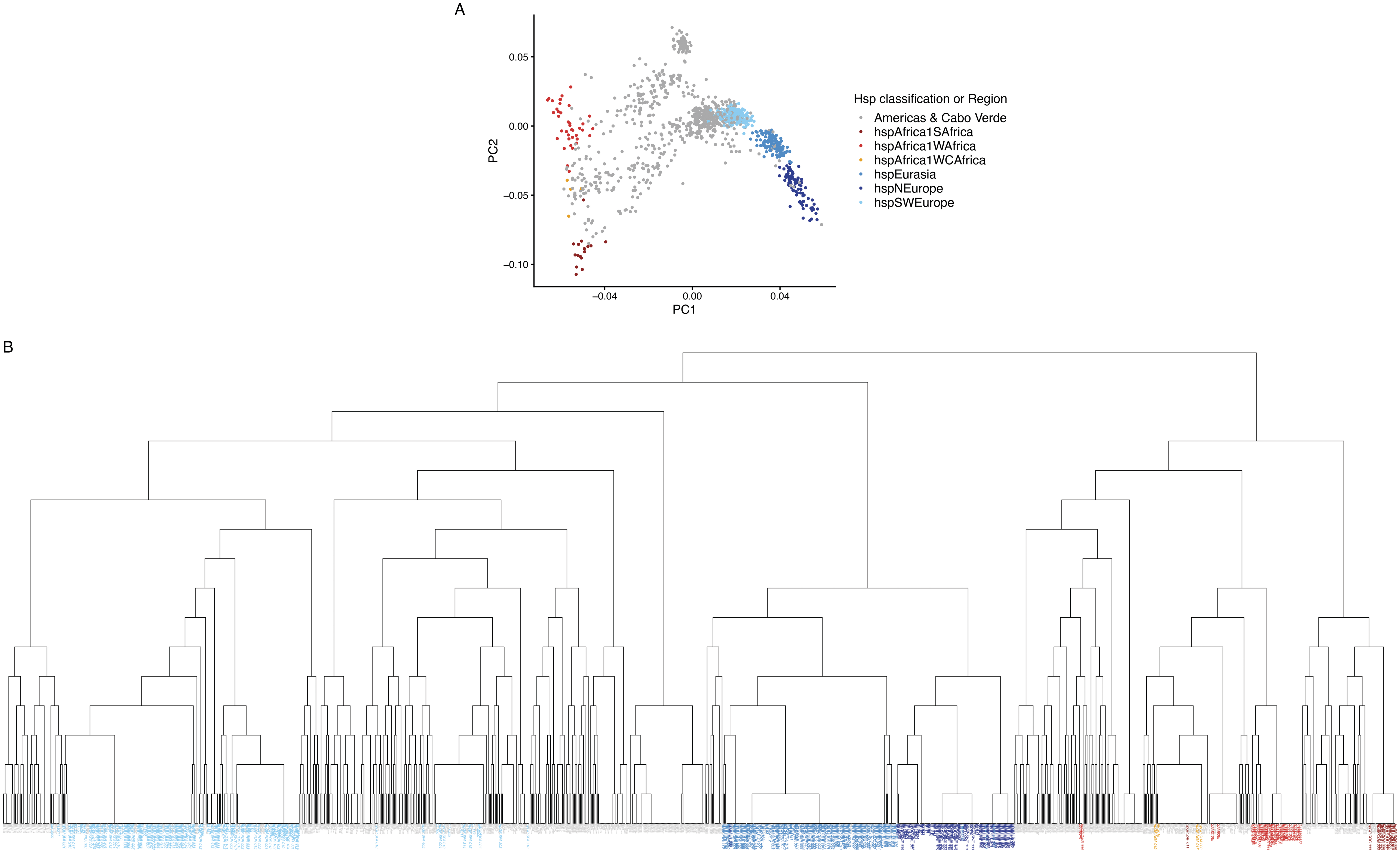

### elife_Figure1-figure_suppl7_revised.pdf

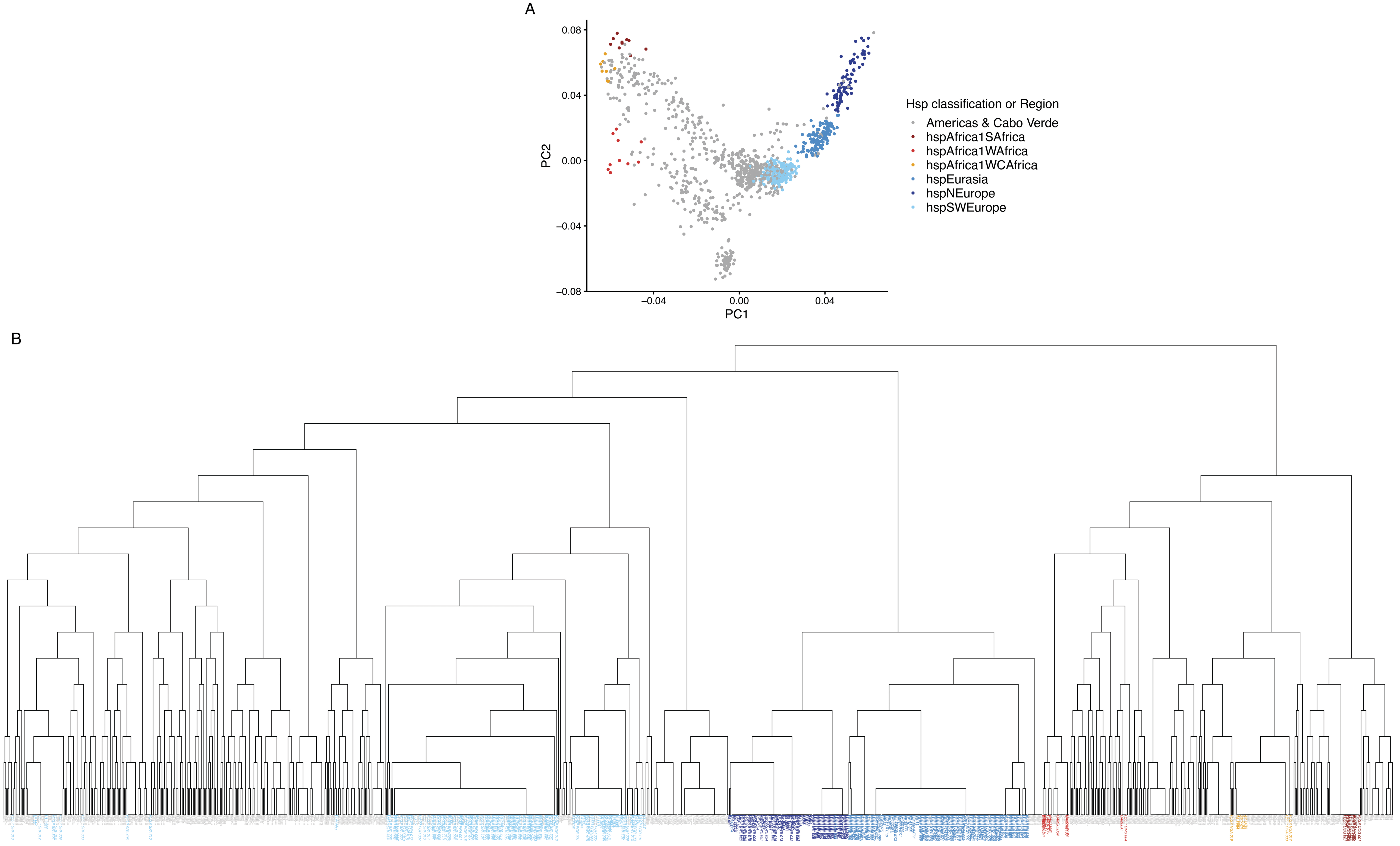

### elife_Figure1-figure_suppl8_revised.pdf

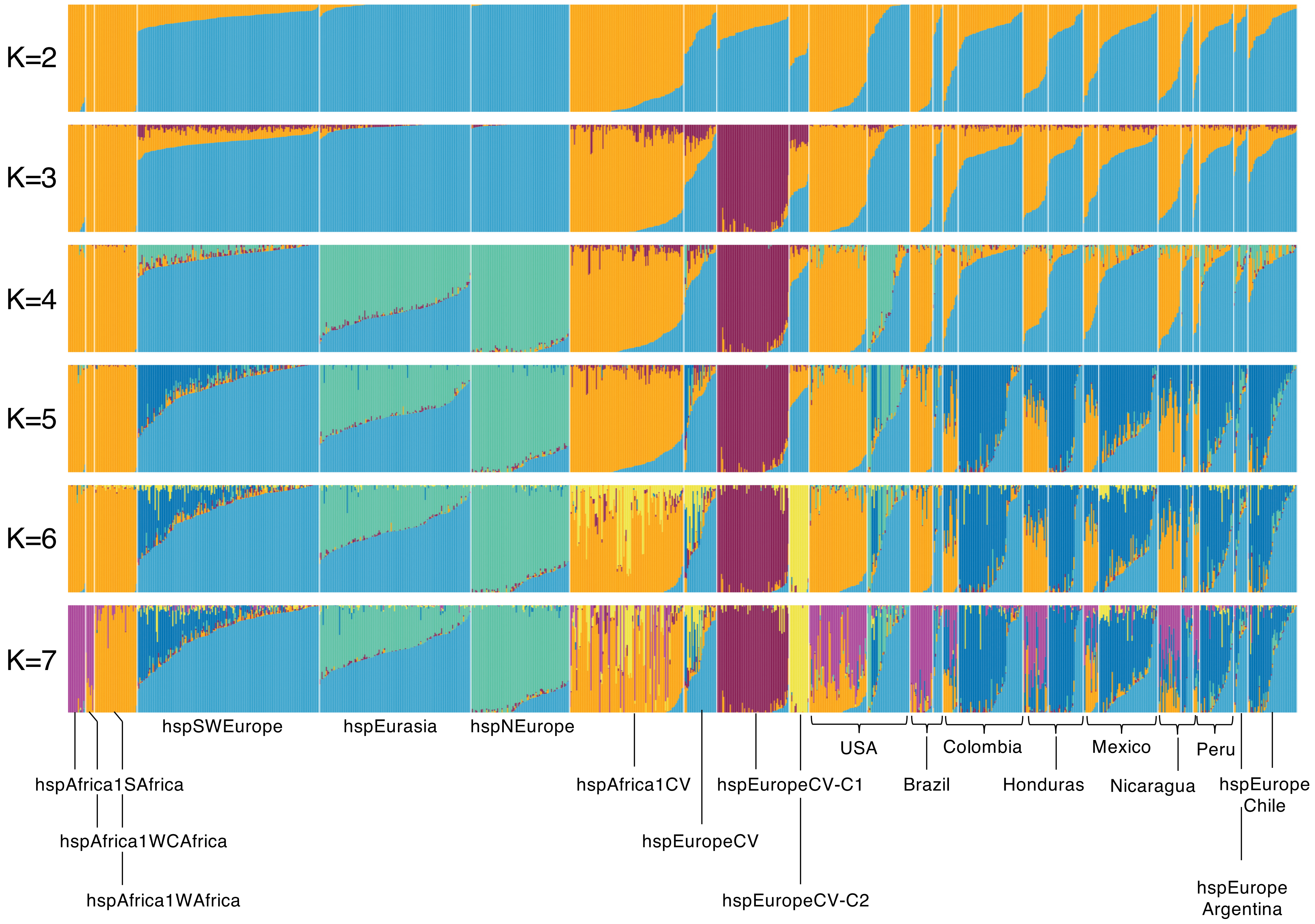

### elife_Figure1-figure_suppl9_revised.pdf

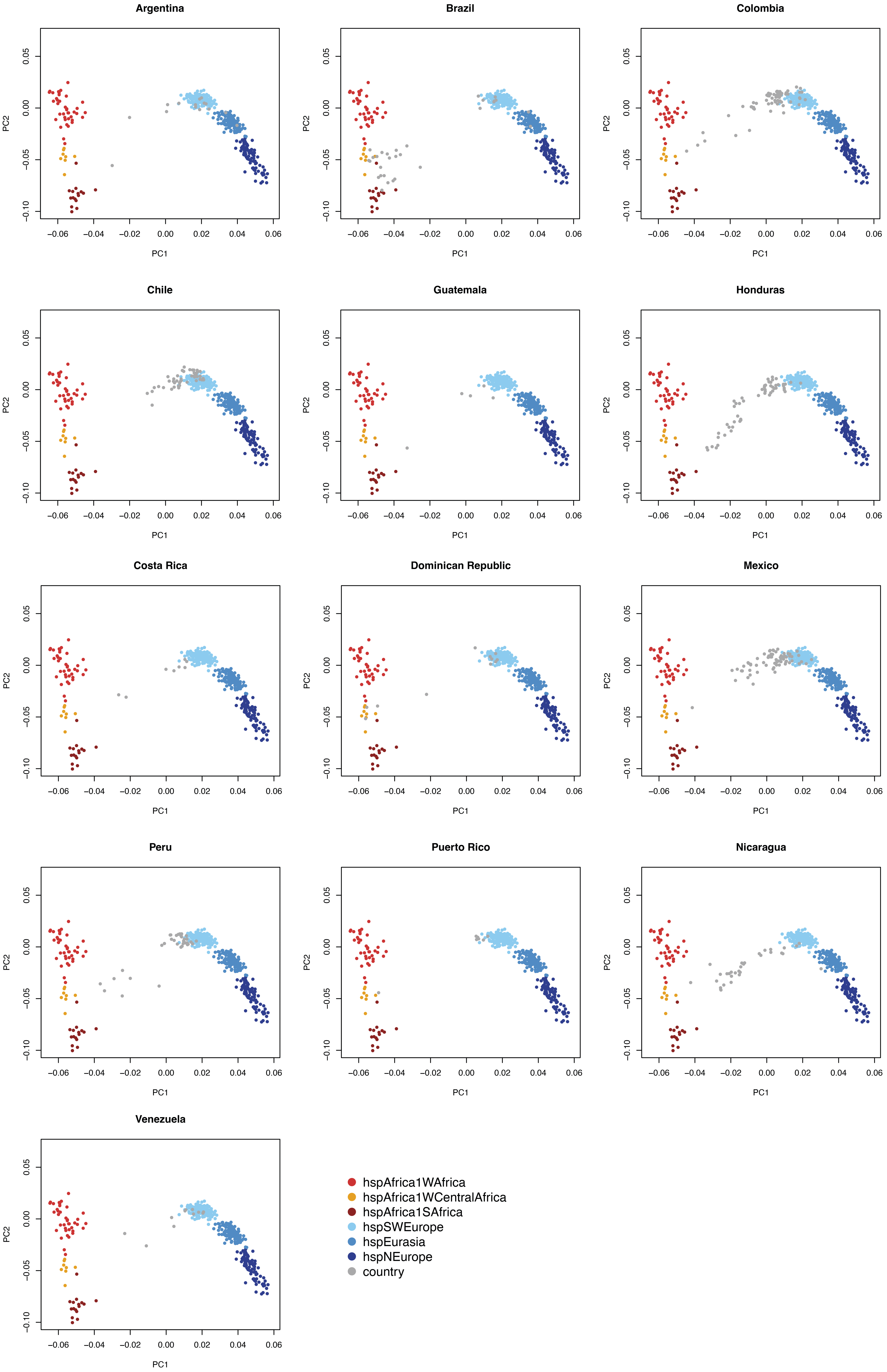

### elife_Figure2-figure_suppl1_revised.pdf

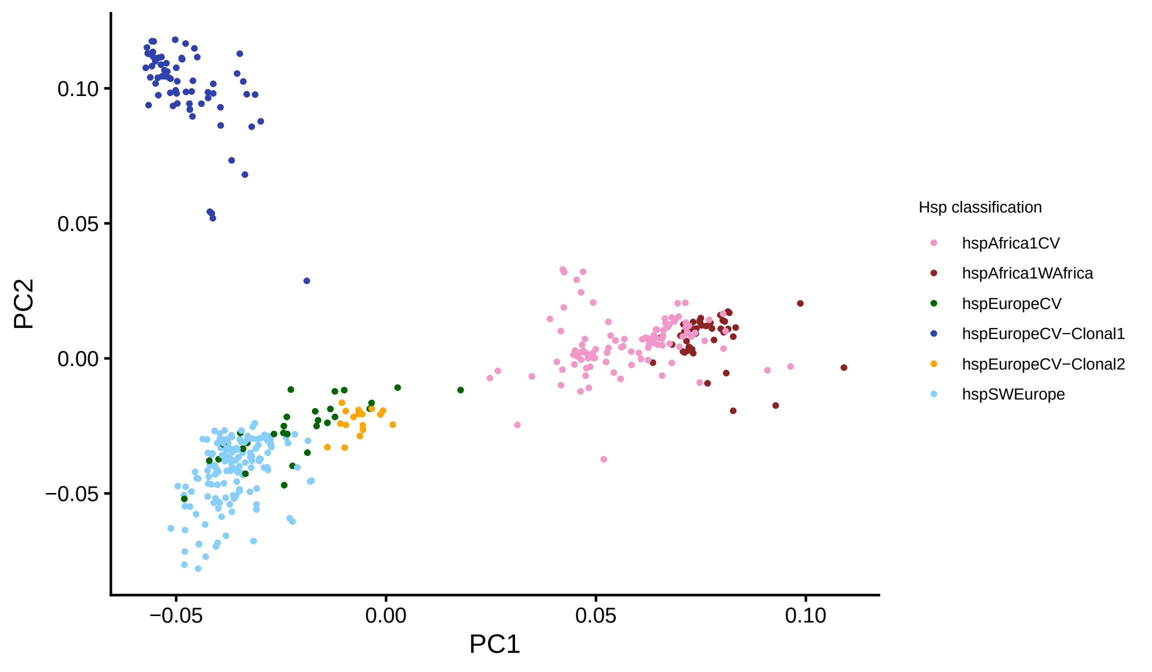

### elife_Figure2-figure_suppl2_revised.pdf

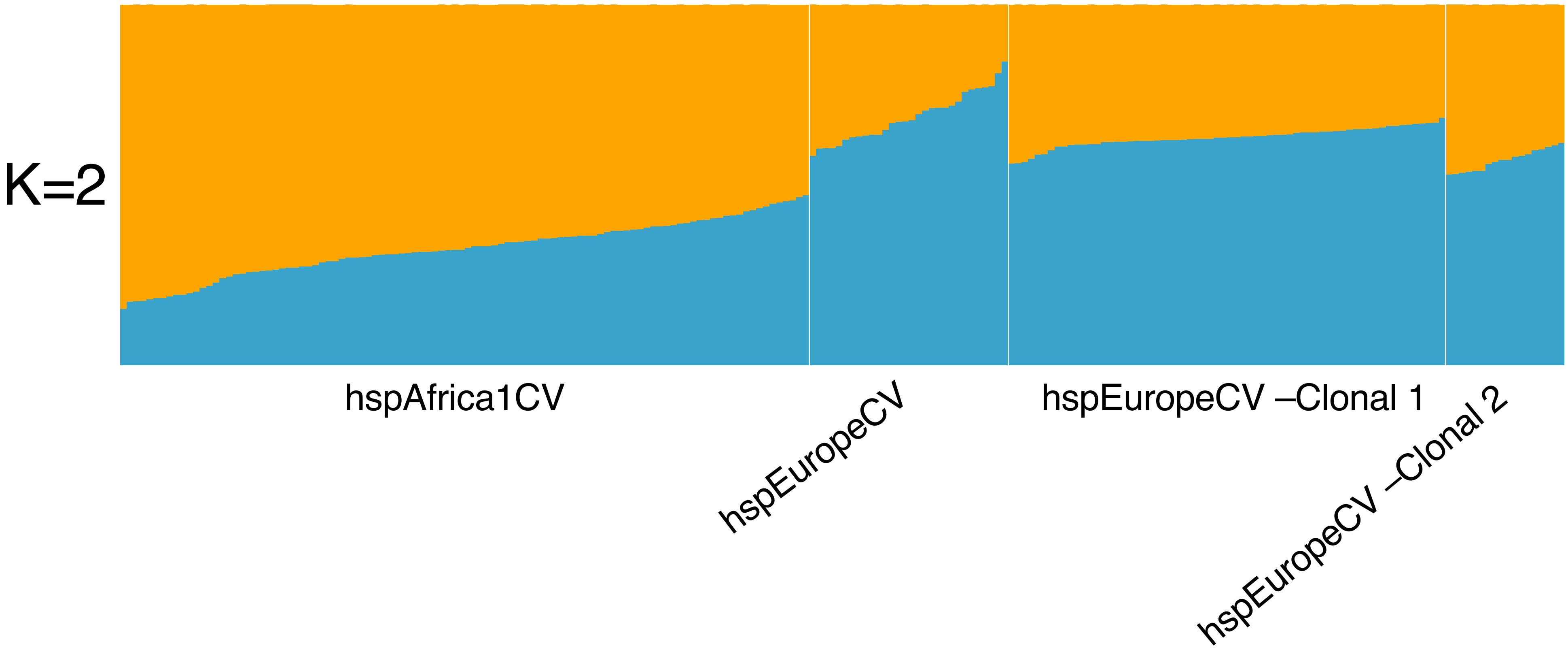

### elife_Figure2-figure_suppl3_revised.pdf

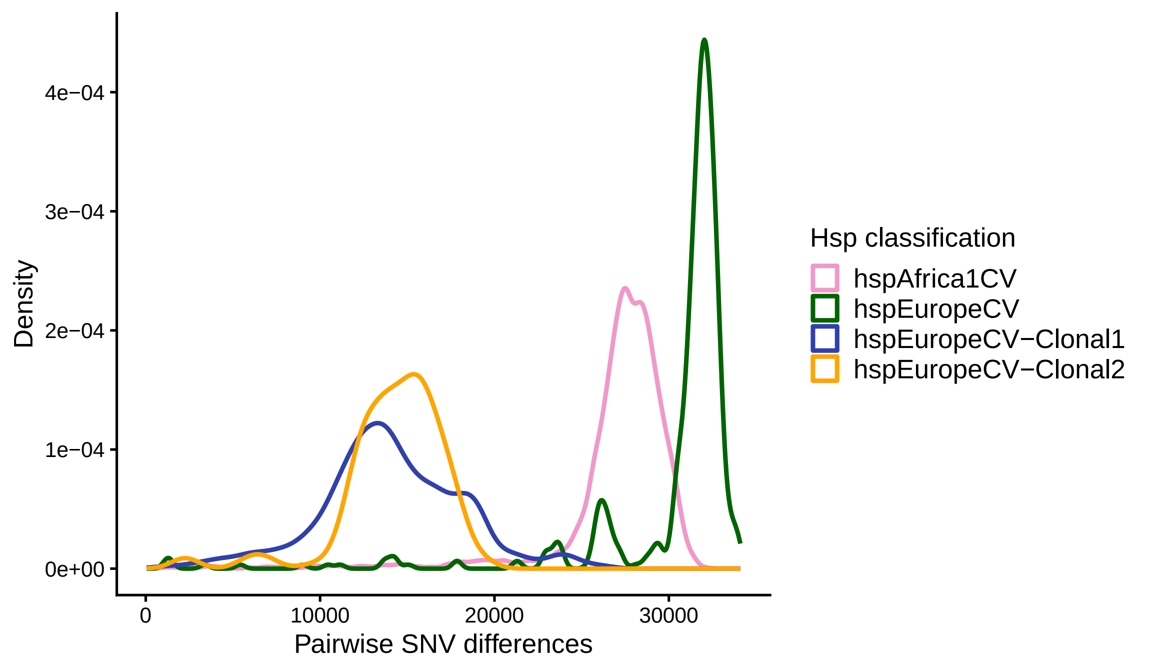

### elife_Figure2-figure_suppl4_revised.pdf

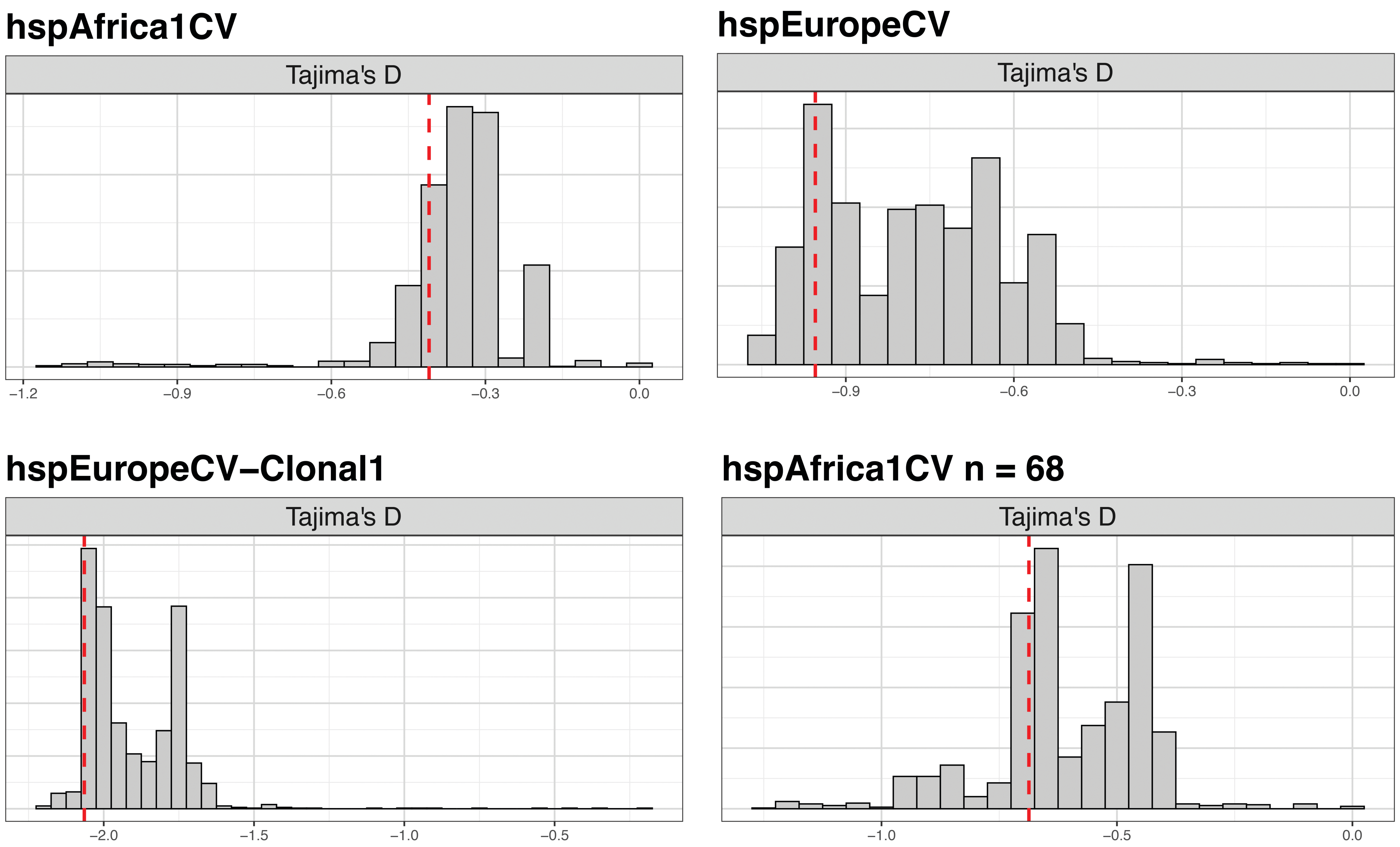

### Table1-Figure_suppl1__revised.pdf

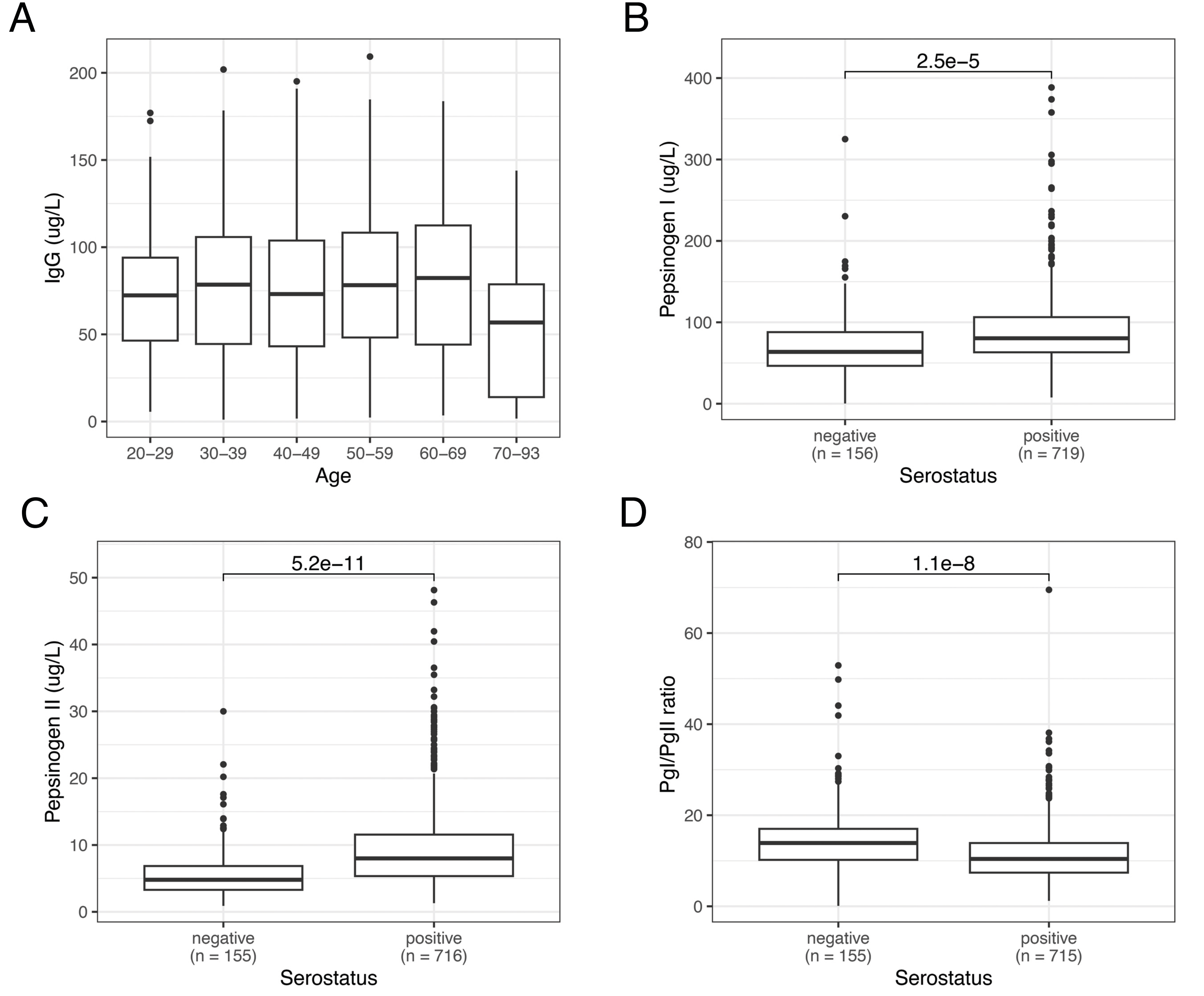
